# Psychosocial experiences modulate asthma-associated genes through gene-environment interactions

**DOI:** 10.1101/2020.07.16.206912

**Authors:** Justyna A. Resztak, Allison K. Farrell, Henriette E. Mair-Meijers, Adnan Alazizi, Xiaoquan Wen, Derek E. Wildman, Samuele Zilioli, Richard B. Slatcher, Roger Pique-Regi, Francesca Luca

## Abstract

Social interactions and the overall psychosocial environment have a demonstrated impact on health, particularly for people living in disadvantaged urban areas. Here we investigated the effect of psychosocial experiences on gene expression in peripheral blood immune cells of children with asthma in Metro Detroit. Using RNA-sequencing and a new machine learning approach, we identified transcriptional signatures of 20 variables including psychosocial factors, blood cell composition and asthma symptoms. Importantly, we found 174 genes associated with asthma that are regulated by psychosocial factors, and 349 significant gene-environment interactions for gene expression levels. These results demonstrate that immune gene expression mediates the link between negative psychosocial experiences and asthma risk.

## Introduction

Psychosocial experiences have long been recognized to affect human health(Miller et al. 2009). Intrapersonal processes (e.g. emotionality(Pressman et al. 2019; Smith et al. 2004), interpersonal social relationships(Repetti et al. 2002; Robles et al. 2014)) and broader structural environments (e.g. neighborhood quality and socioeconomic status (SES)(Gallo and Matthews 2003) are all associated with the morbidity and severity of diseases such as asthma(Harrison 1998), cancer(Meyer and Mark 1995), cardiovascular disease (Everson-Rose and Lewis 2005), as well as mortality rates(Holt-Lunstad et al. 2010; Chida and Steptoe 2008). Asthma is a chronic inflammatory disease of the respiratory tract that disproportionately affects children(Moorman et al. 2012). It is one of the costliest pediatric health conditions(Weiss et al. 2000), and a leading cause of school absenteeism(Akinbami and Centers for Disease Control and Prevention National Center for Health Statistics 2006). Financially struggling cities, such as Detroit, are at an especially high risk for asthma morbidity and mortality(Sullivan et al. 2002). While environmental and genetic factors lead to the development of asthma and affect the health of children with asthma(von Mutius 2000, 2009; Umetsu et al. 2002), psychosocial stress is a critical factor contributing to asthma severity(Wright et al. 1998, 2005; Shankardass et al. 2009; Chen and Miller 2007; Sandberg et al. 2000). Understanding the biological pathways underlying these associations is crucial to strengthen the causal claims linking psychosocial experiences and health.

The growing field of social genomics investigates how various dimensions of a person’s social and psychological environment influence gene expression(Cole 2014; Slavich and Cole 2013; Cole 2009; Toyokawa et al. 2012; Galea et al. 2011). There is ample evidence for links between gene expression in blood and three major categories of psychosocial experiences: socioeconomic status (SES)(Chen et al. 2009), social relationships(Robles et al. 2018; Stanton et al. 2017; Powell et al. 2013) and emotionality(Farrell et al. 2018; Segman et al. 2010). Beyond single gene analyses, previous studies in this area(Slavich and Cole 2013; Cole 2014, 2009) identified a pattern of differentially expressed genes referred to as the *conserved transcriptional response to adversity* (CTRA). The CTRA is characterized by increased expression of genes involved in inflammation and decreased expression of genes involved in type I interferon antiviral responses and IgG1 antibody synthesis(Fredrickson et al. 2013). However, these studies investigated a limited set of psychosocial experiences and did not resolve whether these pathways are causally linked to health outcomes or rather a consequence of disease status.

Several approaches have been developed for investigating the role of gene expression in complex trait variation(Nica et al. 2010; Marigorta et al. 2017; Võsa et al. 2018; Nica and Dermitzakis 2008). Recently, Transcriptome Wide Association Studies (TWAS) and other Mendelian randomization (MR) approaches have been used to integrate genetic effects on gene expression and on complex traits to establish causal links between a gene and a phenotype(Gusev et al. 2016). MR approaches have been developed in epidemiology to examine the causal effect of a modifiable exposure on disease without conducting a randomised trial. MR designs use the genotype association with the two variables of interest to control for reverse causation and confounding. Here, we use this type of approach to connect genes with complex traits. Traditionally genes are annotated to association signals in GWAS based on physical proximity. TWAS test for an association between gene expression and complex traits, where gene expression is predicted based on genotypes in the GWAS study and independent eQTL data. Notably this goes beyond physical proximity of GWAS signals to genes, and establishes a putative mechanism linking genetic variants to complex traits through genetic regulation of gene expression. Very few studies of genetic regulation of gene expression (expression quantitative trait loci, eQTL mapping) in humans have included comprehensive information on psychosocial exposures, and no study to date has been able to determine the likelihood of a causal relationship between psychosocial experiences, gene expression and asthma. This study aims at filling this gap by combining genetic and well-characterized psychosocial data from a cohort of children with asthma living in Metro Detroit (Figure 1a).

**Fig. 1.**
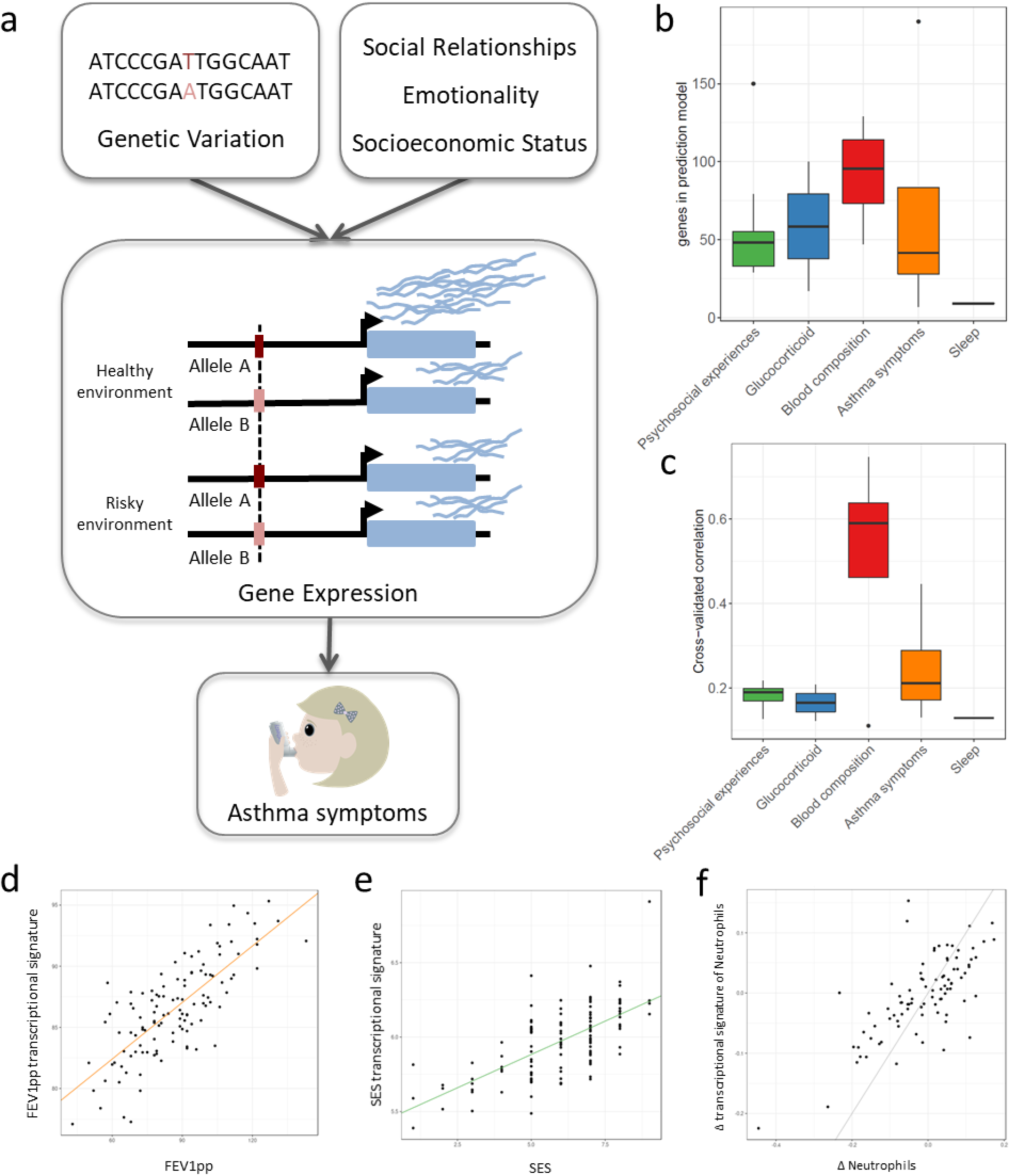
**a** - Central hypothesis, **b** - Number of genes in elastic net regression models that explain at least 1% of variance. Colors represent different categories of variables. **c** - Correlation between cross-validated transcriptional signatures and measured variables for elastic net regression models that explain at least 1% of variance, **d** - Forced Expiratory Volume percent predicted transcriptional signature model fit (Pearson’s rho=0.76, p<0.001), **e** - Socioeconomic status transcriptional signature model fit (Pearson’s rho=0.67, p<0.001), **f** - Longitudinal change in observed Neutrophils (x axis) and longitudinal change in transcriptional signature of Neutrophils (y axis) (Pearson’s rho=0.72, p<0.001, grey=identity line).

The Asthma in the Lives Of Families Today (ALOFT) project was established in 2009 to identify the behavioral and biological pathways through which family social environments impact youth with asthma. This study started during the years leading up to Detroit filing for bankruptcy in 2013 and is still ongoing. Detroit started a marked economic recovery in 2016; yet not all population groups and geographic areas have experienced it simultaneously or to the same extent. To analyze the relationship between psychosocial experiences, asthma and transcriptional regulation, we investigated genome-wide gene expression (RNA-seq) for 251 youth participating in the ALOFT study. For 119 participants, we also collected 53 psychosocial and biological variables (Table S2, Fig. S1). Measures of psychosocial experiences were grouped into five clusters, indicating SES, social relationship functioning, and emotionality. Psychosocial experiences were captured through subjective and objective measures (e.g. negative affect assessed from daily diaries and recorded audio, respectively), as well as global and daily measures.

## Results

### Psychosocial factors and asthma alter the transcriptome

To de-noise and impute psychosocial effects on gene expression for the entire cohort of 251 participants, we developed a new machine learning approach based on generalized linear models with elastic net regularization (GLMnet(Friedman et al. 2010)) and cross-validation. Using this approach we derived transcriptional signatures that represent the portion of the transcriptome that correlates with each psychosocial factor. Analogous methods have been adopted to define transcriptional signatures of T-cell exhaustion in aging(Alpert et al. 2019) and survival in cancer(Asgharzadeh et al. 2006), but have not been previously used for psychosocial factors. We identified significant transcriptional signatures for 32 out of 53 variables (Figure 1b-e, Table S6). We used an independent longitudinal dataset to validate the transcriptional signatures. We considered the changes in the observed variable between two time points (≥1 year) and compared it to the longitudinal changes in the transcriptional signature. Note that the transcriptional signature is imputed for the second time point from gene expression samples that are not included in the training set. We found significant correlations in the observed and imputed changes for the majority of variables (Spearman correlation p-value<0.05; e.g. Fig 1f, Table S7).

Transcriptional signatures of the SES measures showed a strong overlap with each other (Fig. 2a), suggesting that they may have very similar molecular effects or measure the same factors. However, we also saw correlations across all three variable categories. For example, subjective SES was significantly correlated with objective parental responsiveness, family conflict, and self-reported self-disclosure, which is the extent to which the youths talk about their thoughts and feelings (r=-0.26, p=0.004, r=0.25, p=0.006, r=0.53, p=6.8*10^-10^, respectively). Measured psychosocial factors were also associated with interindividual variation in gene expression for several genes. For example, perceived responsiveness and self-disclosure were associated with changes in gene expression for 143 and 3279 genes, respectively (Table S5).

**Fig. 2.**
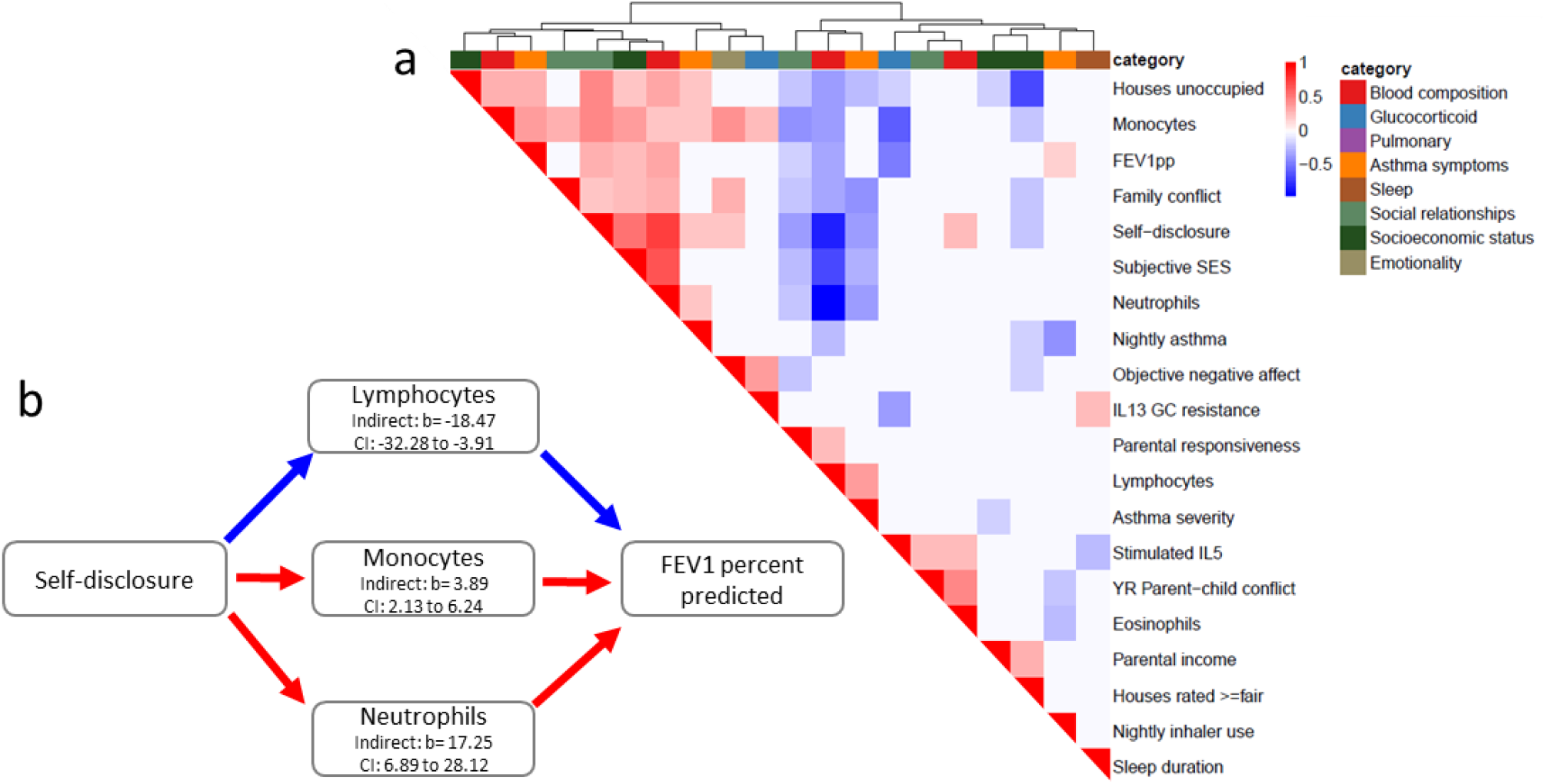
**a** - Heatmap of correlations between transcriptional signatures of variables explaining at least 1% of variance. Heatmap color indicates strength and direction of correlation; white indicates p-value>0.05. Hierarchical clustering of variables is represented above the heatmap, with colors indicating categories for each variable as indicated in the legend. **b** - Mediation analysis between transcriptional signatures of self-disclosure and percent-predicted FEV1, through transcriptional signatures of cell composition.

When we correlated transcriptional signatures of asthma severity with those for psychosocial variables, we observed overlap with SES and social relationships, but not emotionality. In particular, we found significant positive correlations between the transcriptional signatures of lung functioning (percent-predicted FEV1) and psychosocial measures of self-disclosure (r=0.30, p=0.001), objective maternal responsiveness (r=-0.19, p=0.04), subjective SES (r=0.25, p<0.05), and percent unoccupied properties in the neighborhood (r=0.27, p=0.003). These results provide a potential mechanism through gene expression changes in leukocytes for previously reported links between parent-child relationship quality and asthma symptoms(Miller and Wood 2003; Tobin et al. 2015b), as well as SES and asthma symptoms(Litonjua et al. 1999; Mielck et al. 1996). Past research has found emotionality to be a strong predictor of asthma severity(Lehrer et al. 1993). Here we found that the transcriptional signature of self-disclosure was also significantly associated with other measures of asthma, such as nightly asthma symptoms (r=0.22, p=0.02) and asthma severity (r=-0.38, p<0.001), echoing the large body of work on the importance of self-disclosure for health(Pennebaker 1995).

Notably, the transcriptional signatures of blood composition were also associated with asthma symptoms, with a positive correlation for proportion of lymphocytes and negative correlation for proportion of neutrophils (Fig. S7). Given the important role of several blood cell types in asthma severity and exacerbations(Mamessier et al. 2008; Vedel-Krogh et al. 2017; Bigler et al. 2017; Ray and Kolls 2017; Lima-Matos et al. 2018; Casciano et al. 2016; Sinz et al. 2017), we hypothesized that transcriptional changes associated with blood composition mediated the correlations between psychosocial experiences and asthma outcomes. Using mediation analysis, we found significant (p<0.05) paths through all three blood composition signatures, such that, at the molecular level, self-disclosure association with higher pulmonary function could be partially explained by an increase in the proportions of monocytes and neutrophils and reduced proportions of lymphocytes (Figure 2b).

### Genetic interactions with psychosocial factors affect gene regulation

To directly investigate whether transcriptional signatures associated with negative psychosocial experiences contribute to inter-individual variation in asthma risk, we used expression quantitative trait locus (eQTL) mapping combined with transcriptome-wide association analysis (TWAS)(Gusev et al. 2016). TWAS uses eQTLs as instrumental variables to causally link gene expression to phenotypes. To examine local genetic effects on leukocyte gene expression, we performed cis-eQTL mapping and identified 8590 genes (eGenes) with at least one eQTL (10% FDR). These eGenes were enriched in GTEx whole blood eGenes(Aguet et al. 2019) (Fisher’s test OR=3.2, p-value < 2.2*10^-16^), but we also identified additional 1,792 eGenes that were not detected by GTEx in whole blood. These newly-identified genetic effects may be due to limited power in eQTL studies and/or differences between our cohort and GTEx samples in cell type composition, ancestry, age, psychosocial environment, and/or the asthma status.

We used the method for Probabilistic TWAS analysis (PTWAS)(Zhang et al.), which improves upon previous TWAS methods by ensuring only strong instrumental variables are used, and is designed to allow for validating the causality assumption (see Methods). We identified 2,806 eGenes in the GTEx dataset that were causally associated with asthma and allergic diseases (hay fever, eczema and allergic rhinitis) (5% FDR). Of these, 443 were eGenes in our dataset. Here we interrogated whether these causal genetic effects can be modulated by psychosocial factors through gene-environment interactions. To examine the genotype-by-environment effects of psychosocial experiences and blood composition on gene expression, we used the imputed transcriptional signatures for the entire cohort of 251 individuals. In addition to augmenting our sample size, we argue that these transcriptional signatures may better capture the environmental effects on the state of the cells at the molecular level (i.e., after denoising), compared to the observed variables. This is because the observed variable has high levels of noise, and the measured value may not reflect the true biological effect. Therefore, we used the predicted values for all participants, including those for whom the variable was directly measured (denoising). This is similar to the context eQTL approach(Zhernakova et al. 2017) that uses other genes as a proxy variable for the environment, but here the “context” is more easily interpretable because it is defined by a transcriptional signature associated with a specific psychosocial factor. Similarly, cell type composition imputed from gene expression was used to map cell type interaction QTLs for 43 cell type-tissue combinations in the GTEx v8 dataset(Kim-Hellmuth et al. 2019).

For each of the eGenes identified in our dataset, we tested the lead eQTL for an interaction effect (see Methods) with any of the transcriptional signatures. We discovered 349 significant interaction eQTLs across 136 unique genes (10% FDR; Figure 3a, File S7). We found interaction eQTLs for all four blood composition signatures (proportion of lymphocytes, neutrophils, monocytes and eosinophils with 83, 65, 27 and 25 GxE interactions, respectively), which represent cell-type-specific eQTLs (57% of all GxE eQTLs). 93 of the 109 blood-interacting eGenes (85%) were also identified as genes with interaction eQTLs with cell type composition in GTEx whole blood(Kim-Hellmuth et al. 2019). We identified 126 GxE interaction effects on gene expression with psychosocial experiences across 78 genes, including self-disclosure (48 genes), subjective socio-economic status (40 genes), and objective maternal responsiveness (16 genes) (Supplementary File S5). These only partially overlapped (77%) GxE effects observed for blood composition, and included interactions specific to psychosocial factors (Fig. 3c). To evaluate whether the interactions with psychosocial experiences may be mediated by cell composition, we repeated the GxE mapping after correcting for blood cell composition. We observed 130 significant GxE effects (10% FDR) after removing the effect of blood composition differences (Supplementary File S6).

**Fig. 3.**
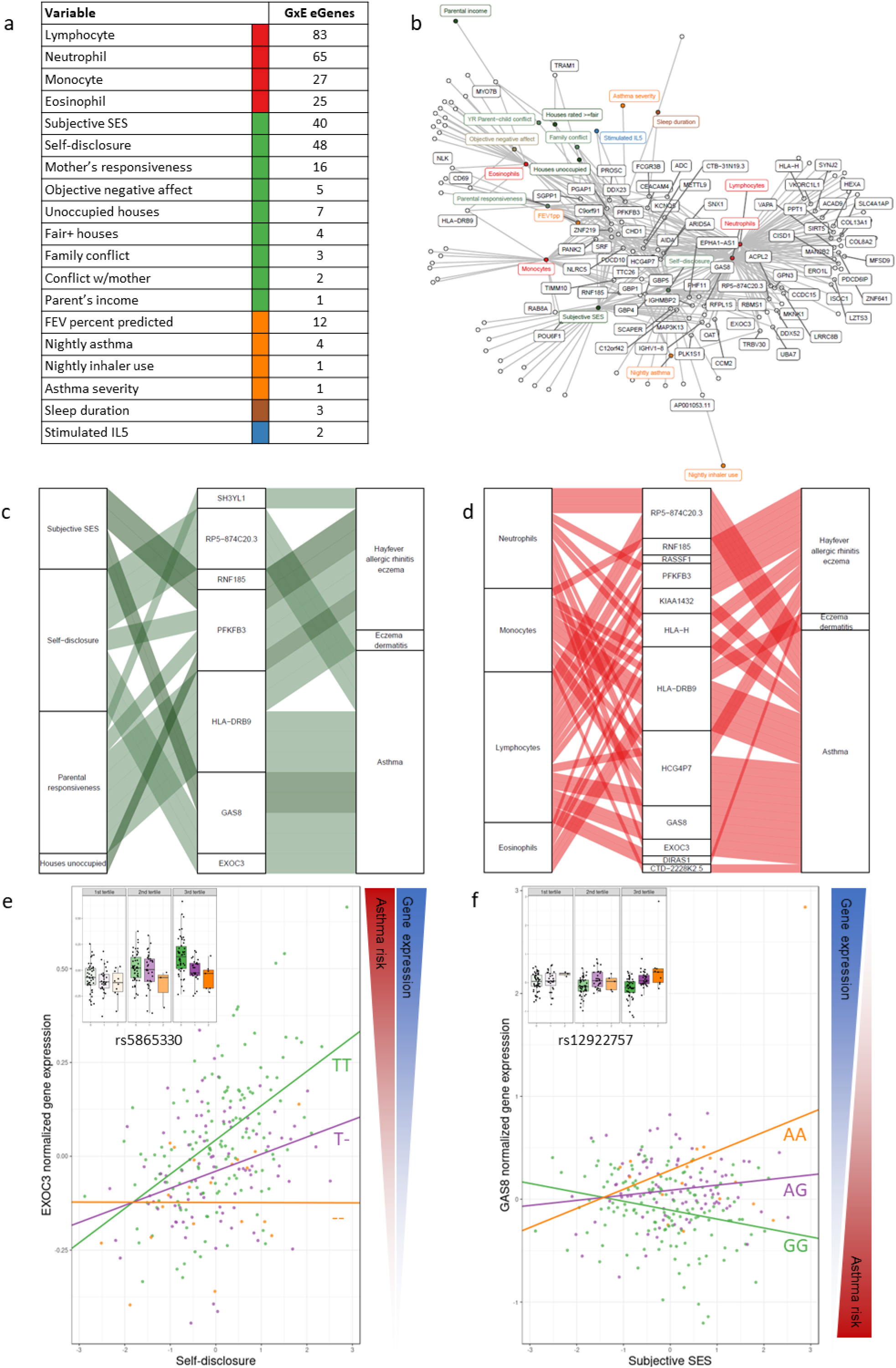
**a** - Number of significant interaction eQTLs (at 10% FDR) for each blood composition trait or psychosocial experience, **b** - Network of interactions between environments and eGenes. Each node represents an eGene with an interaction eQTL (black) or a variable that modulates the genetic effect on gene expression. Only nodes with at least two interactions are labelled. Edges represent significant interaction eQTLS (10% FDR), **c** and **d** - Causal gene-complex trait interactions identified through TWAS are modulated by psychosocial experiences. Psychosocial variables (c) or blood composition (d) are in the left column, eGenes in the central column and complex traits in the right column. A connecting line represents either a causal link between eGene and asthma or allergic disease trait identified through TWAS (middle to right) or a significant interaction eQTL (left to middle). **e** and **f** - Examples of genes causally associated with asthma and with GxE effects that modulate genetic risk. Both genes are causally associated with asthma in TWAS. Each dot is an individual. The boxplot in the inset represents normalized gene expression for the same genes across the three tertiles of the relevant psychosocial variable.

To validate these GxE results we explored the overlap between the GxE genes and previously published datasets that measured interactions with different environments (N=136 genes, see Methods and Supplementary text). We found that 94% of our GxE genes replicated in other datasets of GxE in gene expression (p<0.05). For example, 63 interaction eGenes for psychosocial experiences overlapped with interaction eGenes in response to pathogens(Lee et al. 2014; Barreiro et al. 2012; Çalışkan et al. 2015; Nédélec et al. 2016). This result may indicate that negative psychosocial experiences lead to genotype-specific adverse health effects by influencing the same immune pathways activated by infections. Furthermore, psychosocial experiences may modify the individual response to pathogens and affect health outcomes.

### Risk for asthma is modulated by GxE

We then investigated whether the risk for asthma is modulated through psychosocial experiences (E), and/or GxE effects. Among the genes causally linked to asthma or allergic disease risk by PTWAS, expression of 174 genes was modulated by psychosocial environments, including self-disclosure (124 genes), SES (104 genes), family conflict (30 genes), percent unoccupied houses in the neighborhood (27 genes), parent’s income (19 genes), maternal responsiveness (17 genes), negative affect (e.g. feeling sad or angry, 11 genes), child-reported conflict with parent (9 genes) and percent ≥fair houses in the neighborhood (2 genes) (File S7). The genetic effect on gene expression is modulated by psychosocial factors through GxE for seven genes causally implicated in asthma (4 genes) and allergic diseases (4 genes) (Figs. 3cdef, S11-12). For example, higher expression of the Exocyst Complex Component 3 gene *(EXOC3)* is associated with an increased risk of asthma. We found that self-disclosure, which is the extent to which the youths talk about their thoughts and feelings, increases expression of this gene only for individuals carrying at least one copy of the T allele at rs5865330 (Fig. 3e). The genetic effect was even more pronounced in the highest tertile of self-disclosure (Fig. 3e inset). Lower expression of the Growth Arrest Specific 8 gene *(GAS8)* is associated with an increased risk of asthma. The A allele at rs12922757 increases expression of this gene only in individuals with perceived high socio-economic status, thus reducing the risk of disease (Fig. 3f). A similar effect, and in the same direction is found for *GAS8* and higher self-disclosure (Fig S11-12).

## Discussion

In this study we collected a unique dataset with genome-wide gene expression paired with extensive and accurate assessment of each participant’s biological and psychosocial functioning, across a variety of domains known or likely to be relevant for asthma. We developed a new approach to de-noise and impute the transcriptional signatures of asthma symptoms and psychosocial experiences in peripheral blood. Longitudinal data collected on the same individuals validated the transcriptional signatures imputed on an unobserved later time point, mirroring the changes on phenotype. This demonstrates that the molecular signature of psychosocial experiences on immune cells can track changes over time and can be used to analyze cohorts where these variables are not available.

We showed overlap between transcriptional signatures of asthma symptoms and both socioeconomic status and social relationships, thereby demonstrating that molecular blood gene expression pathways exist through which psychosocial experiences can affect asthma. We further showed that self-disclosure association with higher pulmonary function may be mediated by changes in proportions of monocytes, neutrophils and lymphocytes. Correlations between asthma severity and blood composition are supported by previous findings from the U-BIOPRED cohort where the number of genes differentially expressed between severe asthmatics and healthy controls was reduced by 90% after accounting for blood cell composition(Bigler et al. 2017). This is not surprising as several partially-overlapping endotypes of asthma have been described to date, distinguished by pro-inflammatory contributions from different immune cell types. The most common asthma subtype is characterized by the involvement of T helper type 2 cells (Th2) sensitized primarily to allergens, and subsequent eosinophilic airway inflammation triggered by the type 2 cytokines (particularly Il-5)(Woodruff et al. 2009). However, in non-allergic individuals, eosinophilic inflammation may be triggered by other immune cell types(Brusselle et al. 2013). Elevated levels of neutrophils have been associated with more severe asthma and suggested as an alternative mechanism to eosinophilic inflammation(Ray and Kolls 2017). Further work to dissect the contributions of each cell type can be accomplished in future studies with single cell transcriptomics.

While some of the transcriptional signatures may represent similar underlying processes, the gene expression space that they span cannot be collapsed in one single factor or principal component (Figs. S6 and S7). Some of the signatures could represent factors that can be linked to poverty, while other signatures of social interactions and emotionality are more complex to disentangle. Indeed, the genes sets associated with these variables are enriched for different Gene Ontology functions and pathways. For example, self-disclosure is associated with genes distinctly enriched for neutrophil-mediated immunity (Fig. S5), while parent-reported conflict with child is associated with expression of genes enriched for erythrocyte differentiation. This result suggests that response to negative psychosocial experiences involves processes outside of the scope of the CTRA, which was designed to capture inflammation, antibody production and type I interferon response. Psychosocial experiences change over time because of the children’s development, as a result of broader changes in the urban environments and as a consequence of shifts in family dynamics. These changes of the psychosocial experiences are reflected longitudinally in the gene expression of immune cells and may modify the asthma symptoms and overall health. As we gain additional knowledge on the mechanisms connecting psychosocial experiences to disease risk, these results can be useful to support the need for social interventions that may ultimately lead to improved overall health. The impact of such interventions could be further monitored through gene expression with larger longitudinal samples in future studies.

One outstanding challenge in human complex trait genetics focuses on the portability of polygenic risk scores across population groups or environments(Mostafavi et al. 2020). Pioneering work by our group and others have shown that environmental effects on gene expression and their interactions with genetic factors can play a very important role in regulating genes that are associated with disease(Moyerbrailean et al. 2016; Richards et al. 2019; Findley et al. 2019; Knowles et al. 2017; Nédélec et al. 2016; Zhernakova et al. 2017). This is also applicable to asthma genetics(Rava et al. 2015) and exposure to rhinovirus infection(Çalışkan et al. 2015) and cytokines(Thompson et al. 2020).

Our study demonstrates that many psychosocial experiences leave an impact on gene expression, including genes that are known to be associated with asthma. Furthermore, gene by environment interactions are also found, highlighting that this could be an important factor when evaluating a polygenic risk score for asthma. For example, the individual contribution of a gene can be modified by SES in one direction, and in a different direction for a different gene.

Here, we show that these altered gene expression immune profiles may in turn exacerbate asthma symptoms in children living in inner cities, who are exposed to riskier psychosocial environments. Using human genetics tools we established that psychosocial factors can modulate the causal genetic effects between gene expression and asthma. Importantly, our results demonstrate that psychosocial factors, such as self-disclosure and socio-economic status, modulate genetic risk of asthma and other allergic diseases through altered peripheral blood gene expression.

## Methods

### Study participants

Participants were included from an ongoing longitudinal study, Asthma in the Lives of Families Today (ALOFT; recruited from November 2010-July 2018, Wayne State University Institutional Review Board approval #0412110B3F). The ALOFT study investigates links between family dynamics, biological changes, and asthma morbidity among youth from the Detroit metropolitan area. Participants were recruited from local area hospitals and schools (for recruitment details see Supplemental text). To be included in the study, youth were required to be between 10 and 15 years of age at the time of recruitment and diagnosed with at least mild to persistent asthma by a physician (with diagnosis confirmed from medical records). Youth were screened for medical conditions and medications that might affect asthma and associated biological markers. Only one participant reported current oral corticosteroid use. The full sample included 297 youth and their primary caregivers (typically mothers, referred to as “parent” below). However, only youth with valid gene expression data were included in this investigation. Thus, the sample was comprised of 251 youth (148 boys and 103 girls), whose average age was 12.89 years old (*sd*=1.77 years), and at least one parent. Psychosocial and biological variables, including asthma measures, were available for a subset of 119 participants. For a subset of up to 103 participants we have collected longitudinal data (either 1- or 2-year follow up), which we used to validate the transcriptional signatures. Basic demographic information on these participants is included in Table S1.

### Participant recruitment and collection of psychosocial and biological variables

The parent completed a telephone screening interview to determine eligibility in the study. Written assent and consent were obtained from the participating youth and their parent, respectively.

#### In-lab assessments

The participating youth and parent visited the laboratory, where they completed background questionnaires on a computer and individual interviews assessing stress and asthma management. Parents reported demographics, including their annual income and education level, and completed measures of subjective socioeconomic status (the McArthur ladder (Adler et al. 2000)), neighborhood stress (Ewart and Suchday 2002), conflict with their child (the Parental Environment Questionnaire (Elkins et al. 1997)), and depressive symptoms (the CES-D (Radloff 1977)). The zip code for each family was also collected and used to retrieve objective measures of neighborhood quality based on census block data from 2010 and Data Driven Detroit (collected in 2009), including the percentage of houses rated as fair in quality or better, percentage of houses currently unoccupied, and the percent of people in that area living below the poverty line.

At the same time, youth reported on demographics, warmth received from their mother (Parental Behavior Inventory (Schaefer 1965)), conflict with their mother, the quality of their family environment (the Risky Families Questionnaire (Felitti et al. 1998; Taylor et al. 2004)), depressive symptoms (the Child Depression Inventory (Kovacs 2011)), and the frequency and severity of their asthma symptoms (the Teen Asthma History). Youth also reported on their parents smoking inside the household. However, due to low prevalence as well as uncertainty on whether the parents were present in the household during the four days of data collection, we decided to not use this information in our analyses. They also completed a spirometry test using the nSpire Health KoKo PFT, to obtain the following pulmonary measures: FEV1 percent predicted, FVC percent predicted, FEV1/FVC percent predicted. Also at this visit, the youth and parent were given detailed instructions regarding a four-day daily assessment period. The laboratory visit lasted approximately two hours.

#### In-home assessments

For four days following the laboratory visit (2 weekdays and 2 weekend days), youth and their parent completed daily assessments. Both youth and their parent completed daily diaries each evening about their experiences throughout their day, and sleep diaries each morning about the quality of their sleep. Daily diaries contained items assessing their positive (i.e., happy, interested, excited, and proud) and negative (i.e., sad, angry, upset, worried, distressed) affect, and how much affection and conflict they witnessed between their parents. Youth were also asked to think about the most important and meaningful conversation they had with someone that day, and the extent to which they talked about their thoughts and feelings during that conversation (to measure self-disclosure), and how understanding, validating, and caring their conversation partner was (to measure perceived responsiveness). The sleep diaries contained the Pittsburgh Sleep Scale(Monk et al. 1994), which assesses sleep latency (how long to fall asleep), sleep efficiency (how much time in bed spent sleeping), the number of awakenings throughout the night, the total duration of sleep in hours, and the quality of the sleep. Through the daily and sleep diaries the participants provided information on the following measures of asthma: severity and frequency of daily and nightly asthma symptoms (wheezing, shortness of breath, coughing, chest tightness, other) and nightly inhaler use. Description of daily diary and sleep diary items used in this investigation is included in Supplemental file 1. When youth completed the daily and sleep diaries (i.e., at awakening and before bed), they used a peak flow meter twice to measure peak flow, with the best score between the two assessments used as our measure of morning and evening peak flow. Only daily and sleep diary reports from youth are used in this investigation. Additionally, youth provided four samples of saliva daily for four days at wakeup, 30 minutes after wakeup, before dinner, and immediately before bed using passive drool methods.

Sample time was recorded by participant report, time stamps, and MEMS 6 TrackCap monitors (Aardex Ltd., Switzerland). Samples were initially stored in participants’ refrigerators, but upon return to the lab, saliva samples were stored in the laboratory refrigerator at −20°C until assayed. To reduce positive skewness, we natural log transformed the cortisol values (raw cortisol +1). Hierarchical linear models were run in HLM to extract the average diurnal cortisol intercept, slope, and cortisol awakening response (CAR) for each participant. Finally, participants wore the Electronically Activated Recorder (EAR) in their front pocket or in a belt clip provided from the time they woke up until bedtime. The EAR captured 50 seconds of sound every nine minutes (Mehl et al. 2001). EAR data were coded by trained coders using the Everyday Child Home Observation (ECHO) coding system (Tobin et al. 2015a). Specifically, for this investigation, we use codes of wheezing, positive affect (i.e., happy, interested, excited), negative affect (i.e., sadness, anger, upset, worry, distress), maternal responsiveness (i.e., how much the mother expresses pride, support, and warmth towards the youth), and family conflict (i.e., whether an argument, conflict, fight, or yelling was overheard). Scores for each EAR-observed behavior reflect a mean of the total recordings in which the behavior was observed during waking hours. After completion of the in-home assessment period, the participants returned study materials and the EAR. Youth and parents were compensated for their time.

Additional details on the measures collected in-home are provided in File S1. Descriptive and reliability statistics can be found in Supplemental Table S2. Correlations between measures can be found in Supplemental Figure S1.

#### Biological sample collection

Following the daily assessment period, a peripheral blood draw was conducted for each youth participant. Each youth provided 16, 4 and 8 ml of peripheral blood collected into Vacutainer Cell Preparation Tubes (Becton Dickinson and Co., East Rutherford, NJ) for PBMC (FICOLL gradient vacutainers), DNA (sodium citrate vacutainer, Fisher Scientific catalog #BD-366415) and RNA (EDTA vacutainer) extraction, respectively. Peripheral blood mononuclear cells (PBMCs) were extracted from this sample, as previously described *(66).* All PBMC samples were phenotyped for glucocorticoid (GC) resistance in an established in vitro assay(Marin et al. 2009) measuring the levels of IL-5, IL-13 and IFN-γ in the supernatant (Quantikine ELISA D5000B, D1300B and DIF-50, R&D Systems, Minneapolis, MN). Specifically, PBMCs cultured in RPMI-1640 solution (Life Technologies, Carlsbad, CA) supplemented with 10% FBS (Life Technologies) and 2% HEPES (Sigma-Aldrich, St. Louis, MO) were stimulated for 48 hours with PMA+ionomycin (phorbolmyristate acetate 25 ng/ml, Fisher Scientific, Hanover, IL; ionomycin calcium salt, 1 μg/ml, Sigma-Aldrich, St. Louis, MO) and treated with hydrocortisone (28 nmol/l, Sigma-Aldrich, St. Louis, MO) or vehicle control. GC resistance was calculated as log-fold change of cytokine level in hydrocortisone condition over control and averaged over two replicates. DNA was extracted using DNA Blood Mini Kit (Qiagen, Germantown, MD), and RNA was extracted using LeukoLOCK™ Total RNA Isolation System (Thermo Fisher Scientific, Waltham, MA).

Each of the aforementioned measures were collected annually for a period of two years (three data collection waves) from participants who provided continued informed consent.

### Genotype data

All individuals in this study were genotyped from low-coverage (~0.1X) whole-genome sequencing and imputed to 37.5 M variants using the 1000 Genomes database by Gencove (New York, NY). This data were used for sample quality control (see: Ancestry QC, sex QC and Genotype QC) and to calculate the top three PCs to use as covariates in all statistical analyses.

### Genotype QC

To detect potential sample swaps that may have occurred in sample processing or library preparation, we compared genotypes of RNA and DNA samples from all individuals. We used samtools mpileup function to obtain genotypes from each individual’s RNA-seq bam files for NCBI dbSNP Build 144 variants and kept only variants with more than 40 reads coverage. We used bcftools gtcheck function to compare genotype calls across all biallelic SNPs in all DNA and RNA samples. RNA samples that failed to cluster with their respective DNA sample were repeated (library preparation and sequencing). If the discrepancy was not resolved these samples were excluded from the analysis. A total of 251 samples passed this QC filter. In the end, the pairwise error rate between genotype calls from RNA and their respective DNA samples from the same individual ranged between 0.03 and 0.12. In contrast, the pairwise error rate between all the other unrelated samples ranged from 0.2 to 0.33.

We also used the DNA-derived genotype information to confirm none of the participants were related. We performed Identity-By-Descent (IBD) analysis by Maximum Likelihood Estimation (MLE) using the R package SNPRelate (version 1.16.0). As input we used random 1500 SNPs passing the following criteria: MAF>0.05, missing rate<0.05, LD threshold<0.2. (Fig. S3).

### Ancestry and sex QC

For the 119 individuals for whom the data was available, we plotted self-reported ethnicity against percent global African ancestry defined as the sum of West, East, Central and North African global genetic ancestries calculated by Gencove (Fig. S2a). All samples were in agreement with self-reported ethnicity. Three participants who identified as Multiracial were found to be of admixed African and European ancestry based on genotype analysis provided by Gencove. To check consistency of self-reported sex against genetic data, we plotted fraction of reads mapping to the Y chromosome for all samples. We noted a clear separation between the sexes with no outliers (Fig. S2b).

### RNA-seq data collection and pre-processing

Total RNA was extracted from leukocytes collected on LeukoLOCK (Thermo Fisher) and preserved at −80°C. All RNA samples had a RIN of at least 6 measured on Agilent Bioanalyzer. Library preparation was performed in batches of up to 96 samples (with multiple samples from the same participant always processed within the same batch) on 1-4 ug total RNA, per standard Illumina TruSeq Stranded mRNA library preparation protocol, and sequenced on Illumina NextSeq500 to a depth of 21 million (M) to 76M reads, mean 41M reads (150 bp paired-end). HISAT2(Kim et al. 2015) was used to align demultiplexed reads to the human genome version “GRCh37_snp_tran”, which considers splicing and common genetic variants. Aligned and cleaned (deduplicated) reads were counted using HTSeq and GRCh37.75 transcriptome assembly across 63,677 genes. Post-sequencing quality control included removal of samples with excess PCR duplicate rate (>60%), and genotype QC check against respective DNA sample. For all gene expression analyses, genes on sex chromosomes and genes with expression below 6 reads or 0.1 counts per million in at least 20% of samples were dropped. The final RNA-seq dataset consists of 251 unique samples and 18,904 genes.

### Differential gene expression analysis

We used DESeq2 v1.22.1(Love et al. 2014) to test for differential gene expression across the 23 psychosocial experiences using a likelihood ratio test (LRT) in 119 individuals from the first wave of data collection. To adjust for potential confounders we included as covariates the 3 top PCs of a matrix of possible confounders that included: RIN, site of RNA extraction, library preparation batch, percent reads mapping to exons, percent non-duplicate reads, age, sex, height, weight, top 3 genotype PCs and the four transcriptional signatures of blood composition, Fig. S4). Many of these confounders are very correlated and the 3 top PCs explained 99.7% of their variance. Table S3 represents correlations between individual covariates and the three top PCs of the covariate matrix. In short, PC1-PC3 largely represent weight, height, and age, respectively. For each tested variable, the LRT is then used to compare between two models: GE ~ cvPC1 + cvPC2 + cvPC3 + tested_variable (full model) (full model) and GE ~ cvPC1 + cvPC2 + cvPC3 (reduced model). To control for FDR we used the default independent filtering step and multiple test correction implemented in DESeq2. Table S5 lists differentially expressed genes at 10% FDR; supplementary file S2 contains the full results of the analysis. To compare genes differentially expressed across psychosocial experiences we used PCA analysis using the *irlba* v. 2.3.2 package in R/3.5.2 on z-scores (log2 fold-change/SE (log2 fold-change)) for the top 50 genes with lowest p-value for each tested variable (Fig. S6).

### GO and pathway enrichment analyses

We used the R package *clusterProfiler(Yu et al. 2012)* to run GO, KEGG and REACTOME enrichment analyses (hypergeometric test) across genes upregulated and downregulated compared to the background of all expressed genes (Fig. S5). Enriched categories were defined at 5% FDR.

### Imputation and de-noising of transcriptional signatures

We developed an approach to impute and de-noise a transcriptional signature for psychosocial, environmental and other phenotypic variables based on Generalized Linear Models with Elastic-Net Regularization. First, we normalized the count data using the voom function in the *limma* v3.38.3 package in R(Law et al. 2014). Second, we regressed out the following confounding factors: RIN, percent reads mapping to exons, percent non-duplicate reads, site of RNA extraction, library preparation batch, sample collection wave, age, sex, height, weight, genotype PC1, genotype PC2, genotype PC3. Third, we used the R package *glmnet* v2.0-16 in R/3.5.2 (gaussian model), with a relaxed alpha=0.1 to allow for highly co-regulated genes to be included in the prediction model. Leave-one-out cross-validation was used to evaluate the best models. We used the cross-validated mean square error (MSE) metric, and its standard deviation to evaluate which signatures were more predictive. We calculated the R^2^ for each of the models based on the % MSE reduction from cross validation. To compare the results that would be achievable with the CTRA-based approach, we used the same method but we limited the molecular signature to only include the 53 genes that are used to calculate the CTRA score(Fredrickson et al. 2013). 48 of the 53 genes comprising the CTRA are measurable in our sample (CTRA genes below detection: IL1A, IFIT1L, IFITM5, IFNB1, IGLL3). We compared the fraction of variance explained between the CTRA-based and unrestricted models (Fig. S8).

### Longitudinal Replication

We collected a second time-point (approximately one or two-years after the time point used in current analyses) for a subset of 14 variables: parental income, subjective SES, self-disclosure, YR Parent-child conflict, stimulated IL5, IL13 GC resistance, Eosinophils, Lymphocytes, Monocytes, Neutrophils, FEV1 percent predicted, nightly asthma symptoms, nightly inhaler use and asthma severity, to validate the transcriptional signatures. We considered the longitudinal changes in the transcriptional signatures imputed from the new gene expression data and compared them to the changes in the observed variable between the two time points. Note that the transcriptional signature is imputed for the second time point from gene expression samples that are not included in the training set. We used Spearman correlation to compare the changes from the imputed transcriptional signature to those directly observed.

### Mediation analyses

To test whether transcriptional changes associated with blood composition (Proportion of Basophils, Lymphocytes, Monocytes and Neutrophils) mediated the observed correlations between psychosocial experiences (self-disclosure) and asthma outcomes (percent-predicted FEV), we used the PROCESS macro in SPSS(Hayes 2013) to conduct indirect effect analyses on transcriptional signatures (basic mediation, Model 4, testing the indirect path from independent variable X to mediator M to dependent variable Y) using a bootstrapping approach with 20,000 iterations. The indirect path was considered to be significant if the confidence interval produced by this test did not include 0 for 95% of the iterations.

### cis-eQTL mapping

We calculated gene expression residuals (as in the imputation and de-noising approach) and then used FastQTL(Ongen et al. 2016) with adaptive permutations (1,000-10,000). For each gene, we tested all genetic variants within 1 Mb of the transcription start site (TSS) and with cohort minor allele frequency (MAF)>0.1, for a total of 17,679 genes and 82,679,170 variant-gene pairs tested. We optimized the number of gene expression PCs in the model to maximize the number of eGenes. The model that yielded the largest number of eGenes included 18 gene expression PCs (File S4b).

### Interaction eQTL mapping

To identify interaction eQTLs, we considered the lead eQTL for each of the 4943 eGenes identified at 10% FDR by FastQTL (without correcting for gene expression principal components). This is similar to what was done by GTEx(Kim-Hellmuth et al. 2019) and others(Alasoo et al. 2018; Kim-Hellmuth et al. 2017), and equivalent to a very conservative prunning of all SNPs in the entire cis-association region. We did not correct for gene expression principal components because some of them are correlated with cell-composition and the environmental variables, thus complicating the interpretation of the linear model. To reduce impact of potential outliers, we quantile-normalized each transcriptional signature prior to GxE testing. We fit a linear model that includes both the genotype dosage and the marginal environmental effect as well as their interaction: Expression ~ dosage + transcriptional signature + dosage*transcriptional signature. To fit this model we used the lm function in R-3.5.2. We generated an empirical null distribution of 100 million permuted p-values to correct the interaction p-values (Fig. S9). The empirical null distribution was obtained through multiple runs of the model for each tested transcriptional signature-gene pair while permuting the genotype dosages. Storey’s q-value method to control for FDR was applied on the permutation-corrected p-values for all tests within each transcriptional signature separately.

To ensure the signal detected was not solely due to cell composition differences, we repeated the GxE eQTL mapping procedure as above, while correcting for four signatures of cell composition using the following model: Expression ~ Eosinophils + Leukocytes + Monocytes + Neutrophils + dosage + transcriptional signature + dosage*transcriptional signature. Supplementary File S6 contains full results of this analysis.

### Replication analysis of GxE

To validate our GxE results we considered the following GxE studies for which full interaction testing results are available(Barreiro et al. 2012; Lee et al. 2014; Çalışkan et al. 2015; Nédélec et al. 2016; Moyerbrailean et al. 2016; Kim-Hellmuth et al. 2019). We show numbers of our GxE eGenes (FDR<10%) which replicated in other studies (p-value<0.05).

### TWAS analyses

To directly investigate whether discovered effects on gene expression and GxE interactions may contribute to asthma, allergic disease risk and/or behavioral phenotypes, we used PTWAS results(Zhang et al.) (5% FDR) as an independent source of evidence of causality between gene expression levels and asthma/allergic disease risk (File S5). PTWAS utilizes probabilistic eQTL annotations derived from multi-variant Bayesian fine-mapping analysis conferring higher power to detect TWAS associations than existing methods. The evidence for causality from PTWAS is strong for the following reasons: i) we use only strong instrumental variables (IVs) by combining the strength of multiple independent strong eQTLs for each gene and combining information across all tissues; ii) within the PTWAS framework we can then validate the causality assumption for each gene-trait-tissue combination. We found that the exclusion restriction criterion was violated (heterogeneity of independent estimates across multiple strong eQTLs, I^2^ statistic>0.5) in only 0.36% of the gene-trait pairs for which we computed this statistic, none of which overlap our reported results. Using eQTL data across 49 tissues from GTEx v8, we used PTWAS to analyze GWAS summary statistics from several large-scale projects. Here, we specifically focused on the following asthma studies: GABRIEL-Asthma, TAGC-Asthma-EUR, UKB-20002-1111-self-reported-asthma, UKB-6152-8-diagnosed-by-doctor-Asthma, and allergic disease studies: EAGLE-Eczema, UKB-20002-1452-self-reported-eczema- or-dermatitis, UKB-6152-9-diagnosed-by-doctor-Hayfever-allergic-rhinitis-or-eczema.

Additionally we considered other phenotypes that may be relevant for our cohort: chronotype (Jones-et-al-2016-Chronotype, UKB-1180-Morning-or-evening-person-chronotype), sleep duration (Jones-et-al-2016-SleepDuration, UKB-1160-Sleep-duration), and depressive symptoms (SSGAC-Depressive-Symptoms). To identify eGenes in asthmatic children that are causally associated with asthma, we considered all 4,943 eGenes that were used for the interaction eQTL analysis with a significant (10% FDR) marginal effect of the psychosocial experiences from the linear model that includes both the genotype dosage and the marginal environmental effect as well as their interaction: Expression ~ dosage + transcriptional signature + dosage*transcriptional signature.

## Data access

The data are being submitted to dbGAP. Accession number is pending.

## Acknowledgments

We thank Luis Barreiro and Noah Snyder-Mackler for comments on an earlier version of this manuscript, and members of the Luca, Pique-Regi, Zilioli and Slatcher labs for helpful discussions. We thank the study participants and their families for taking part in this study.

This work was supported by a grant from NIH NHLBI (R01HL114097).

## Disclosure declaration

The authors declare no competing interests.

## Author contributions

J.A.R. drafted the work, contributed to the acquisition, analysis, and interpretation of data, contributed to the creation of new software used in the work; A.K.F. substantially revised the work and contributed to the analysis and interpretation of data; H.E.M. contributed to the acquisition of data; A.A. contributed to the acquisition of data; X.W. contributed to the analysis and interpretation of data, contributed to the creation of new software used in the work; D.E.W. contributed to the acquisition of data and to the conception and design in the initial stages of this work; S.Z. contributed to the conception and design of the work, contributed to the acquisition and interpretation of data; D.E.W. contributed to the conception and design of the work; R.B.S. contributed to the conception and design of the work, contributed to the acquisition and interpretation of data; R.P. contributed to the conception and design of the work, substantially revised the work, contributed to the analysis and interpretation of data, contributed to the creation of new software used in the work; F.L. contributed to the conception and design of the work, substantially revised the work, contributed to the acquisition, analysis and interpretation of data.

## Materials & Correspondence.

All correspondence and material requests should be addressed to F.L. and R.P.

## Supplementary information and extended data

Materials and Methods

Figures S1-S12

Tables S1-S6

Files S1-S8

## Notes

### Competing Interest Statement

The authors have declared no competing interest.

