## Supplement for "Psychosocial experiences modulate asthma-associated genes through gene-environment interactions"

#### 1. Supplementary figures:

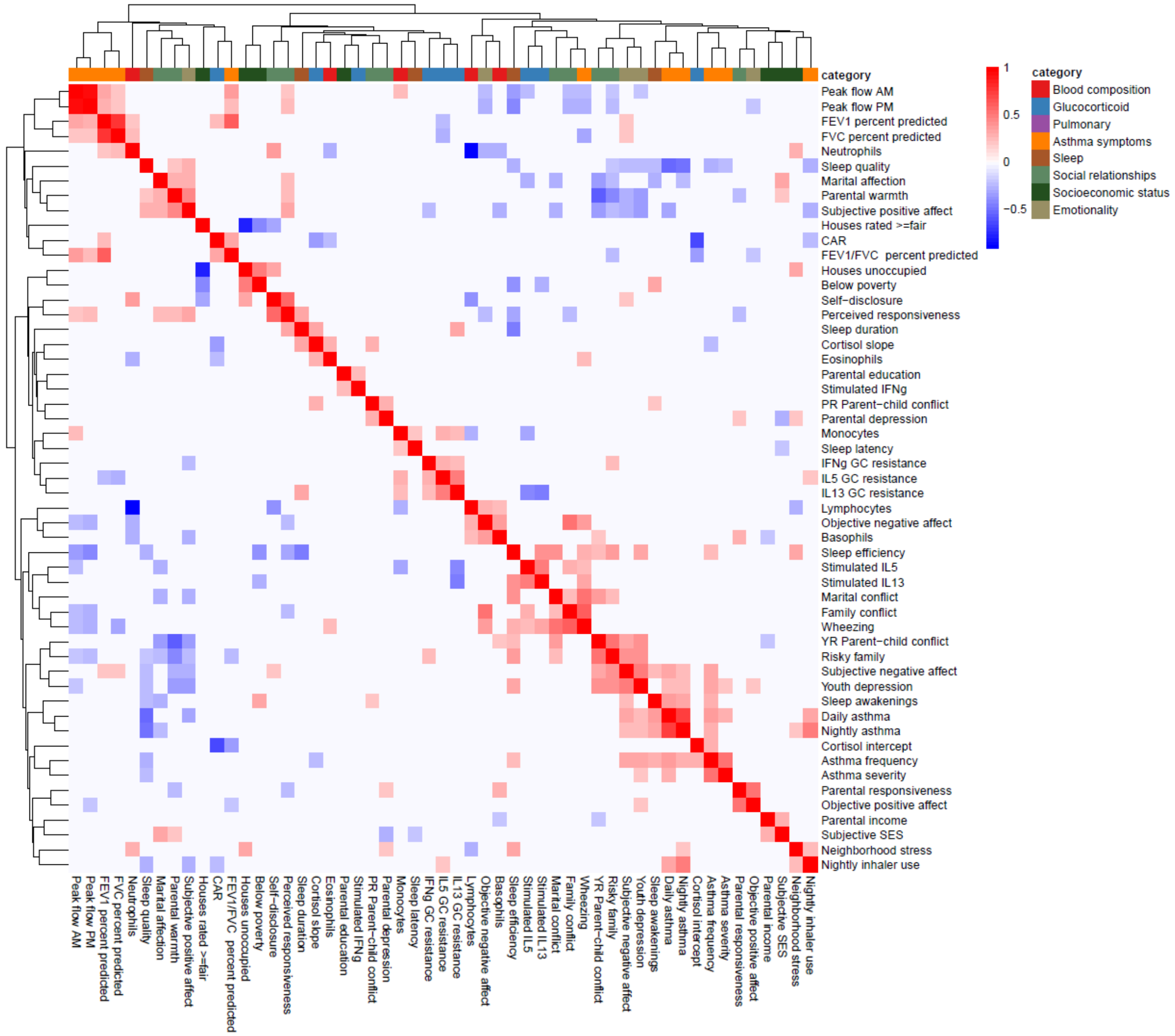


Fig. S1. Clustered heatmap of (Pearson) correlations between all variables used in the study. Color indicates the strength and direction of correlation; white indicates p-value>0.05. Hierarchical clustering is represented on the top and side of the heatmap. Variables are grouped by colors indicating the different categories considered.

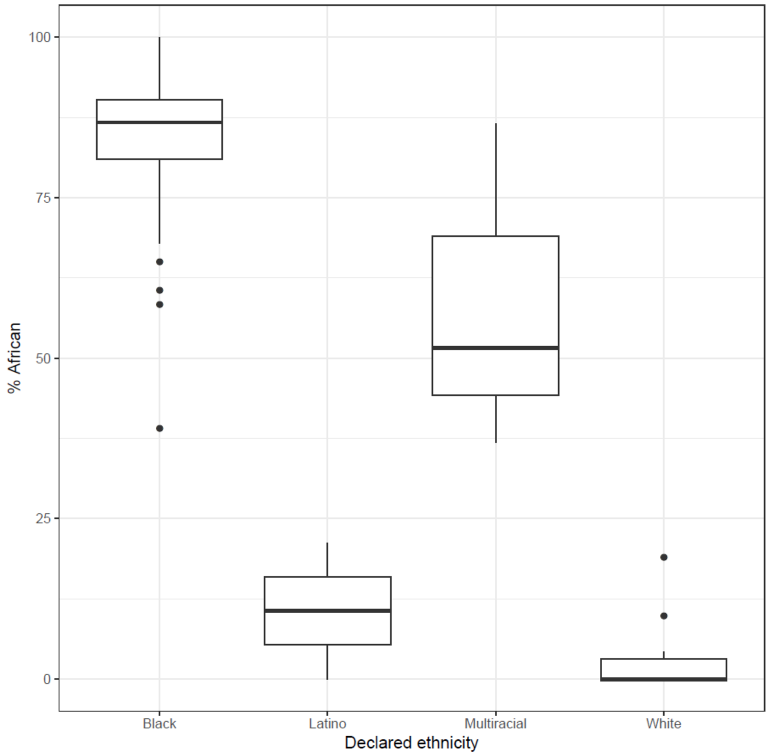


Fig. S2. Self-reported ethnicity (x axis) vs. percent global African ancestry (y axis) in 119 participants for whom declared ethnicity is available.

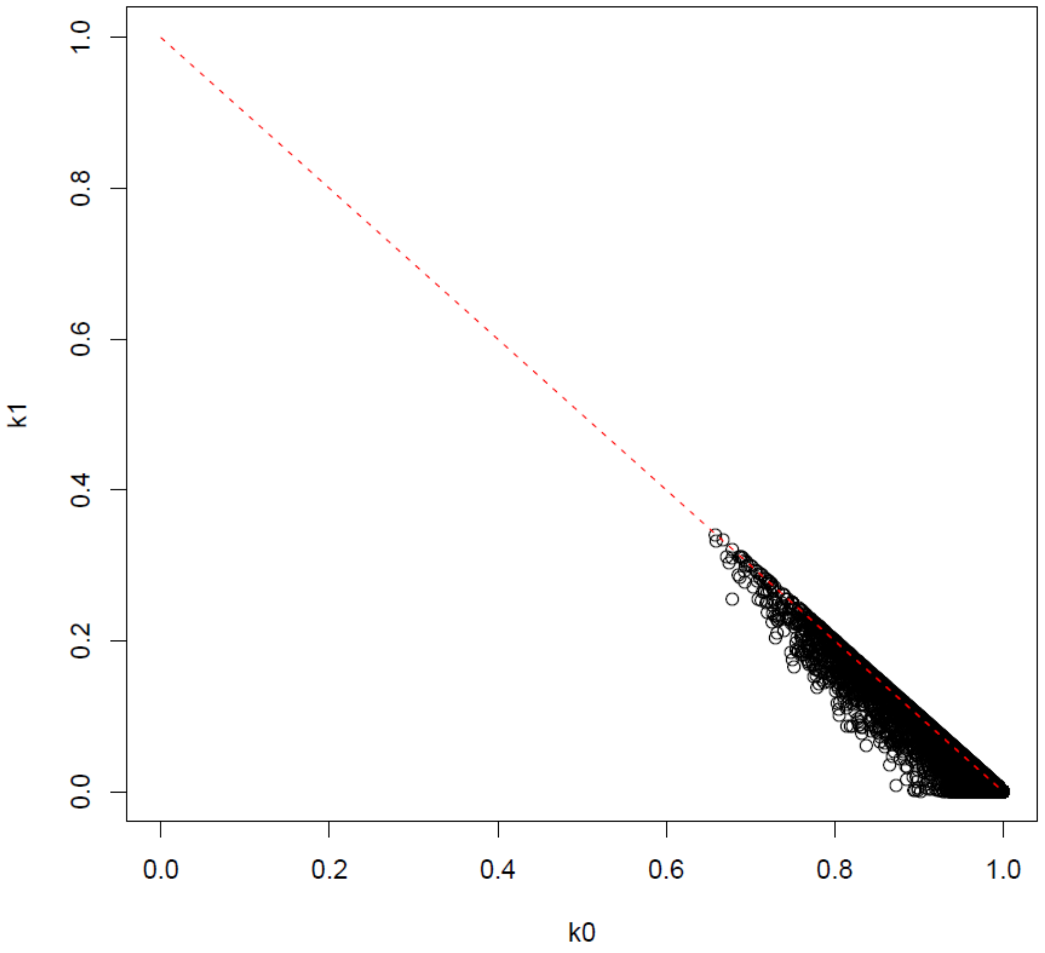


Fig. S3. Result of Identity-By-Descent analysis (MLE) on DNA-derived genotypes for all 251 participants. k0 – probability of sharing zero IBD, k1- - probability of sharing one IBD.

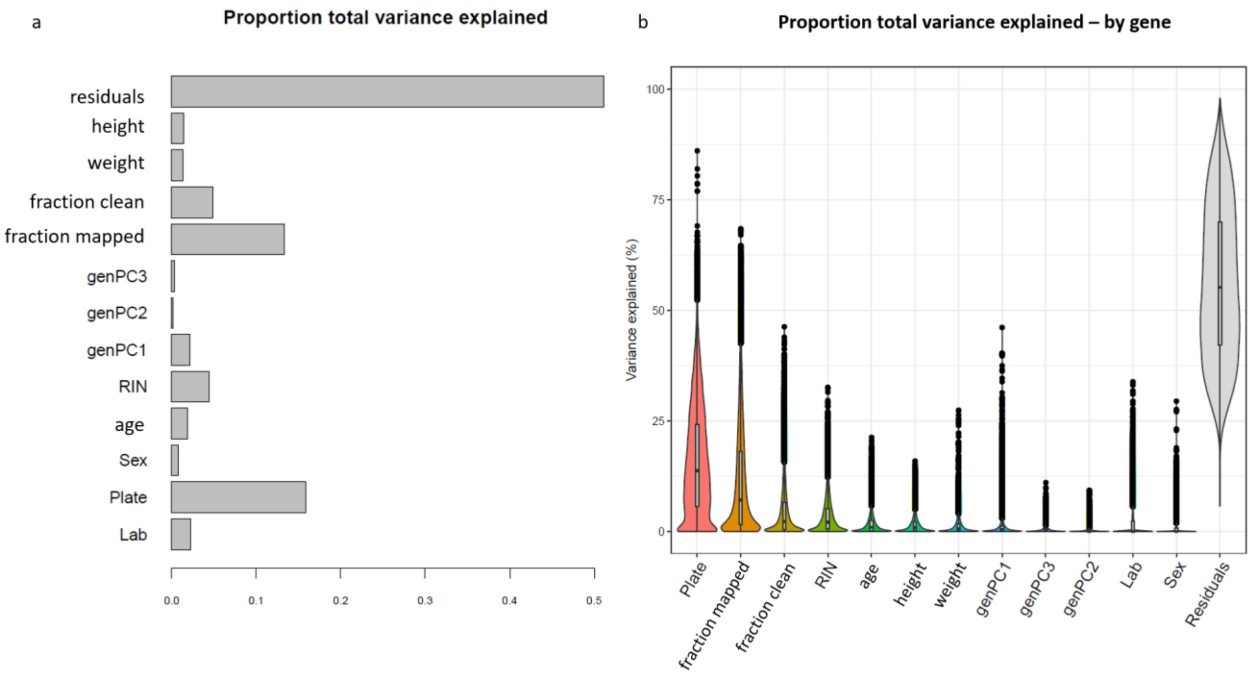


Fig. S4. a - Proportion of global variance in gene expression explained by each covariate tested within a single linear model, b - Proportion of variance explained by each covariate for each analyzed gene.

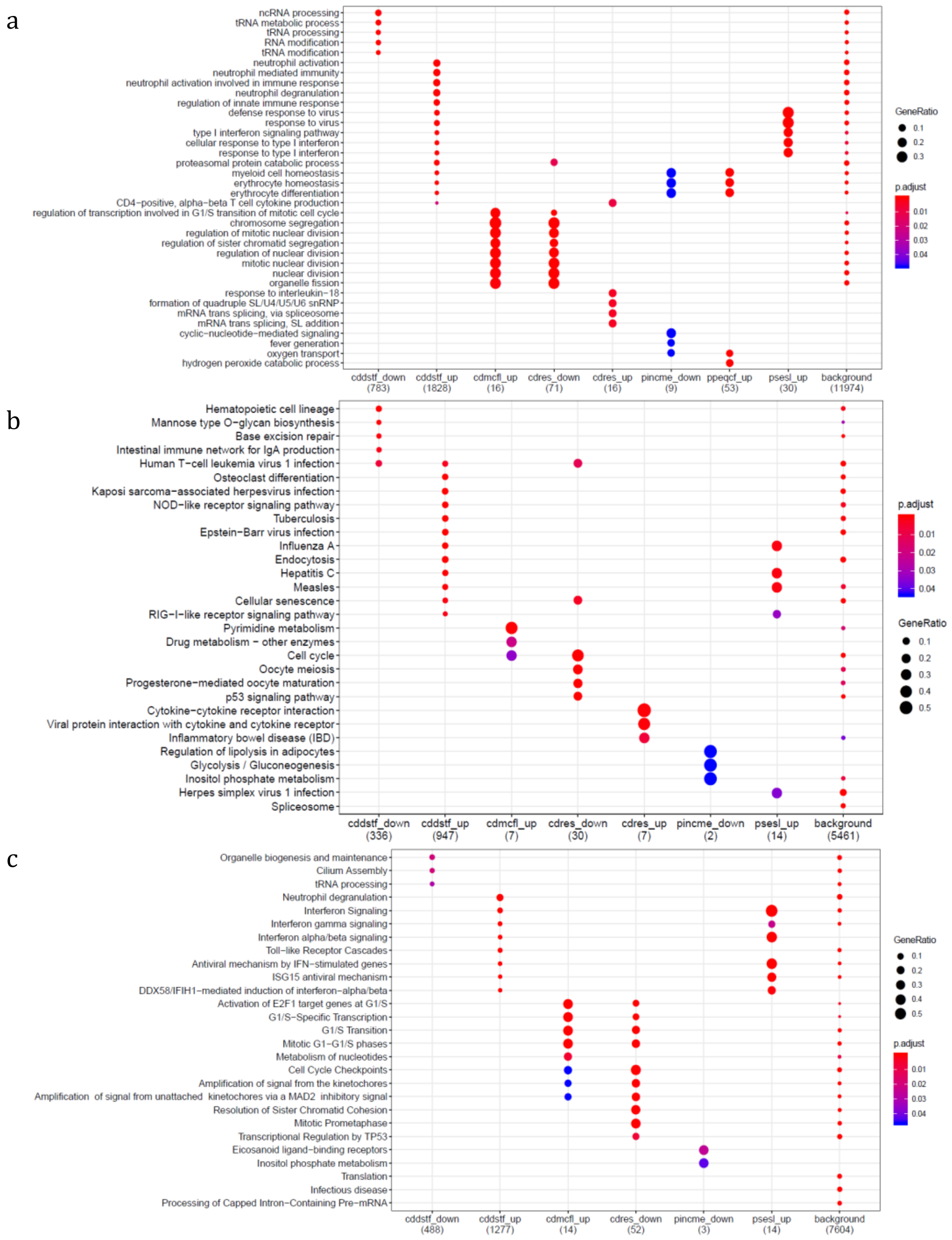


Fig. S5. Gene set enrichment analysis results on genes differentially expressed for psychosocial experiences: a - Gene Ontology, b - Kyoto Encyclopedia of Genes and Genomes, c - Reactome Pathway Database.

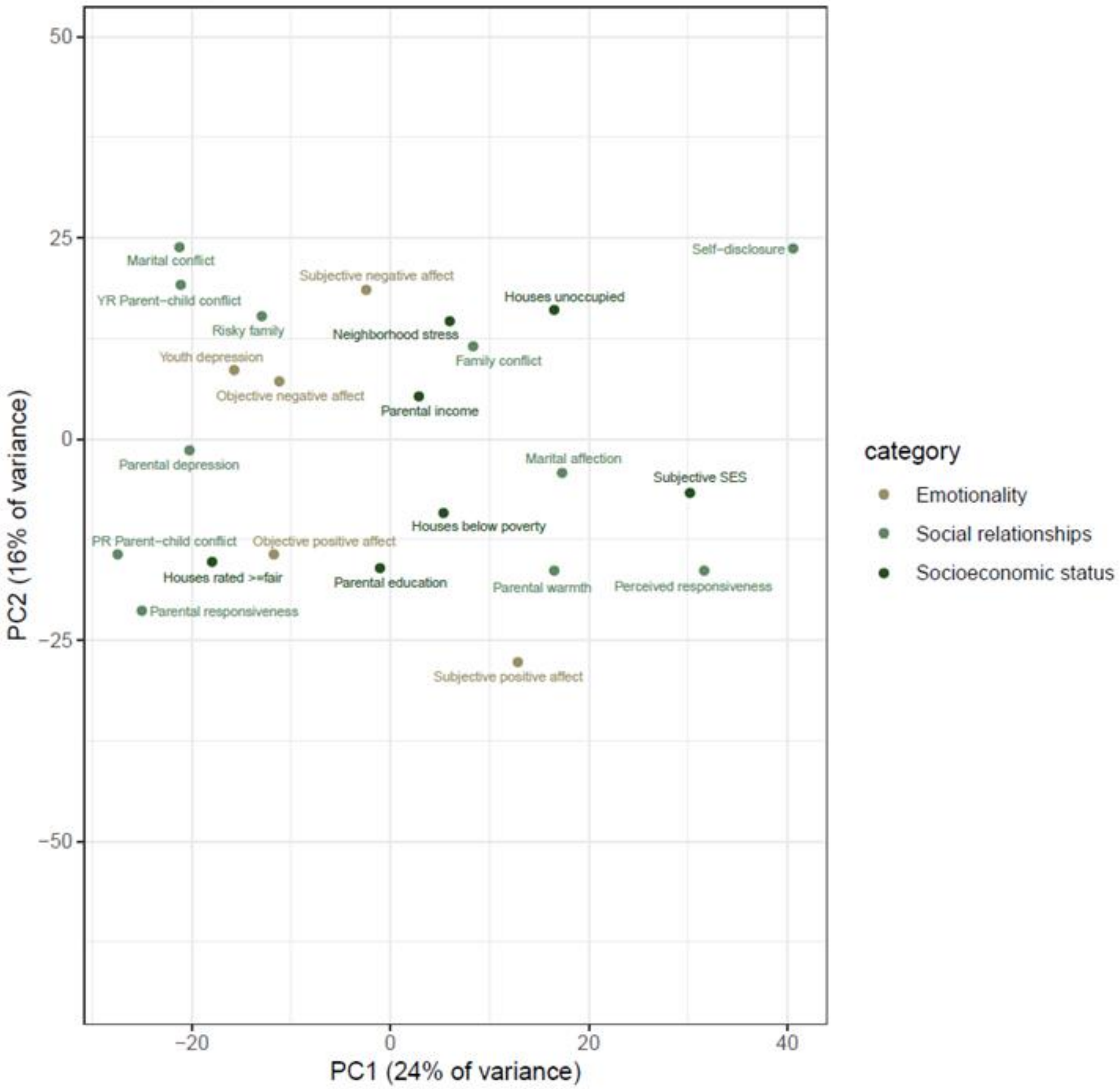


Fig. S6. Principal component analysis of differential gene expression z-scores of the top 50 genes with lowest p-value for each tested variable.

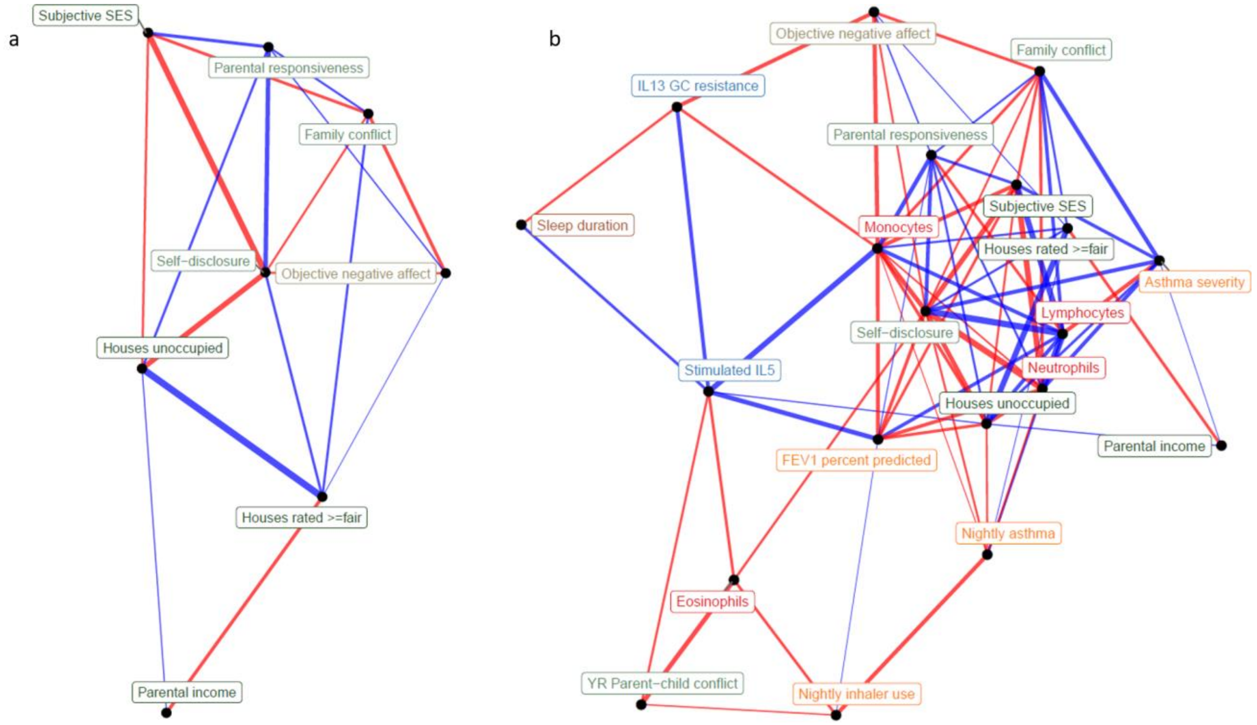


Fig. S7. Network representation of correlations between a - transcriptional signatures of psychosocial experiences, b - all transcriptional signatures (edge width reflects absolute value of Pearson's correlation score, edge color reflects positive (red) or negative (blue) correlation).

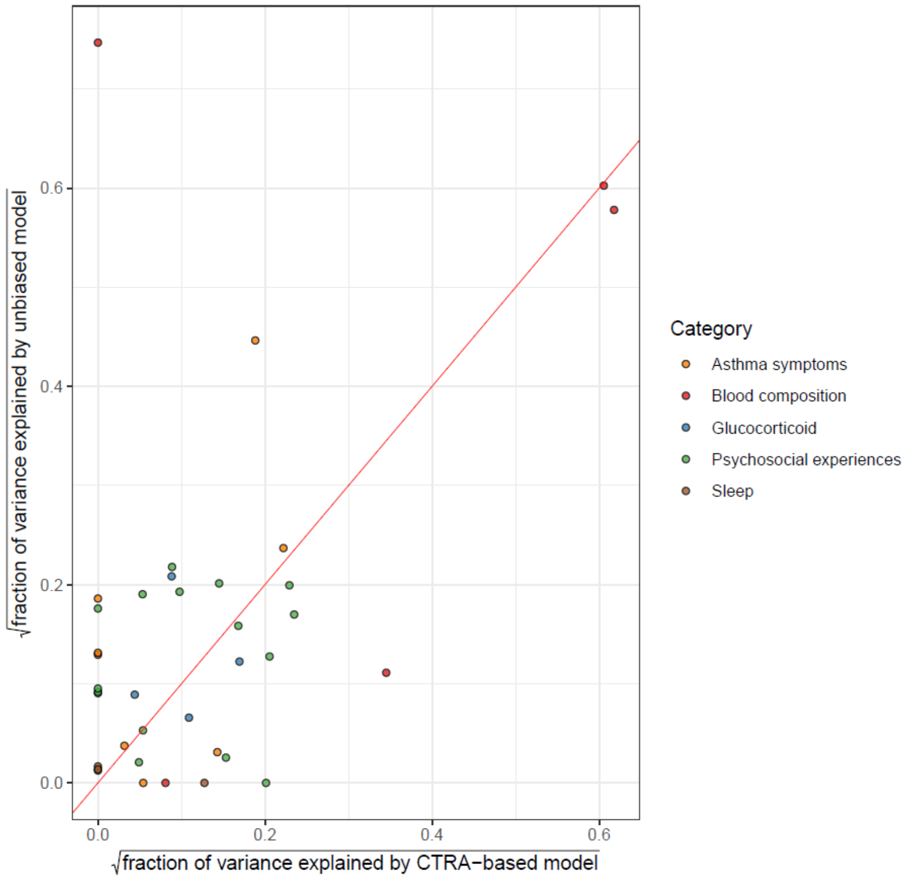


Fig. S8. Comparison of variance explained by CTRA-based (x axis) and unbiased (y axis) elastic net prediction models, color indicates type of variable (red=identity line).

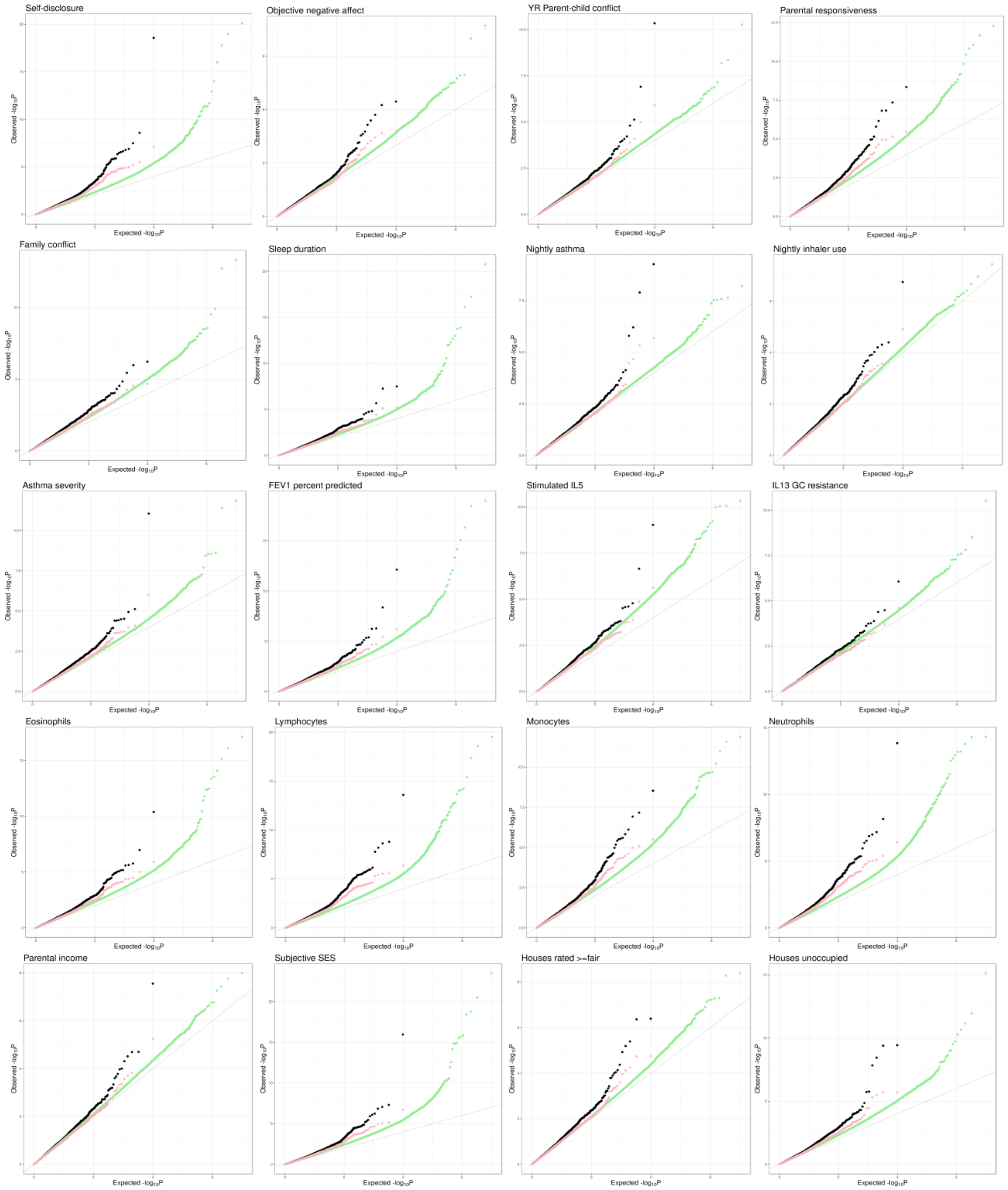


Fig. S9. QQplots of interaction eQTL mapping test p-values (black), permutation p-values (green) and permutation-corrected p-values (pink).

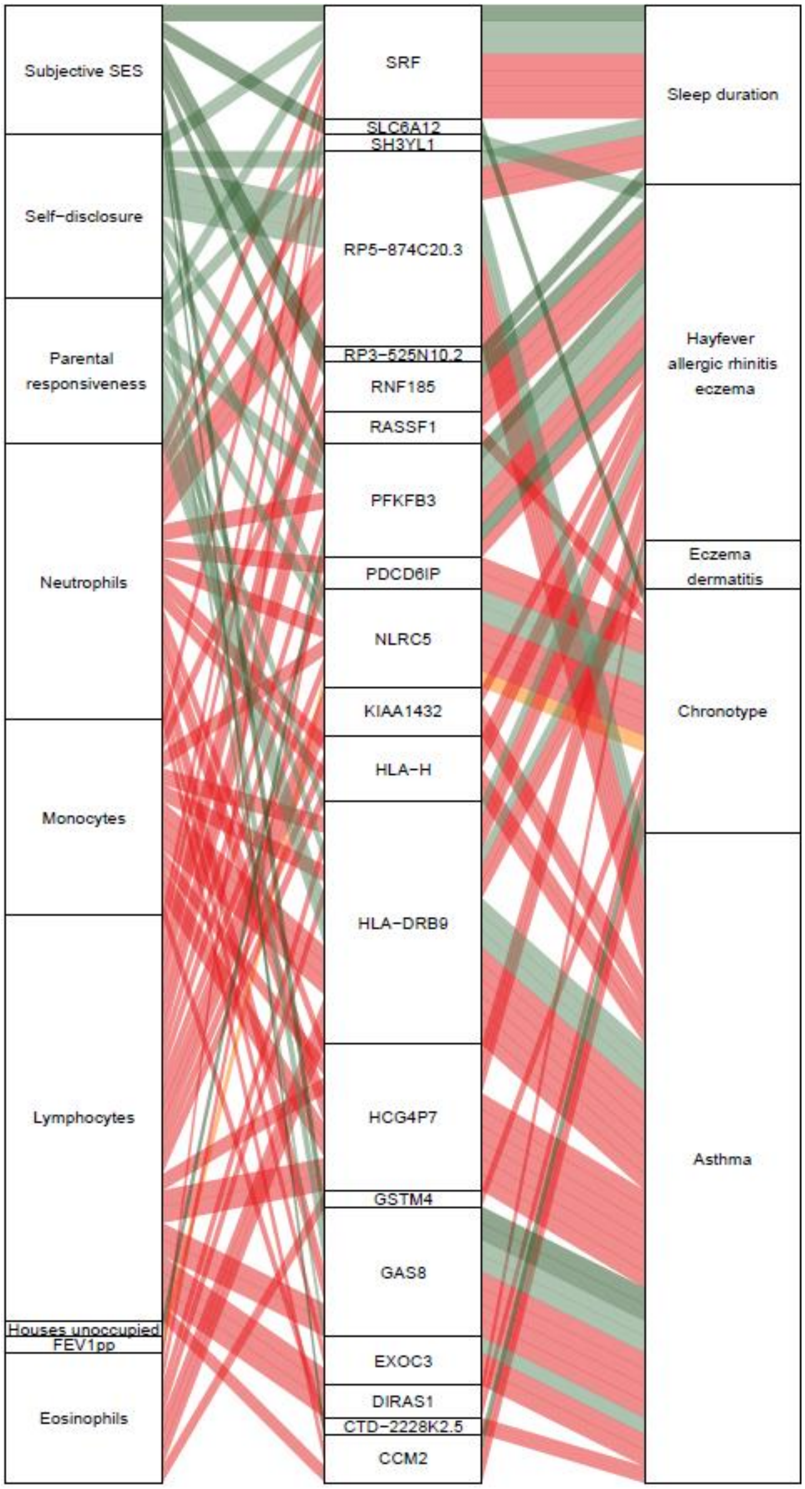


Fig. S10. Causal gene-complex trait interactions identified through TWAS are modulated by psychosocial experiences. Psychosocial and biological variables are in the left column, eGenes in the central column and complex traits in the right column. A connecting line represents either a causal link between eGene and trait identified through TWAS (middle to right) or a significant interaction eQTL (left to middle). Red represents blood composition variables, orange represents asthma variables and green represents psychosocial variables.

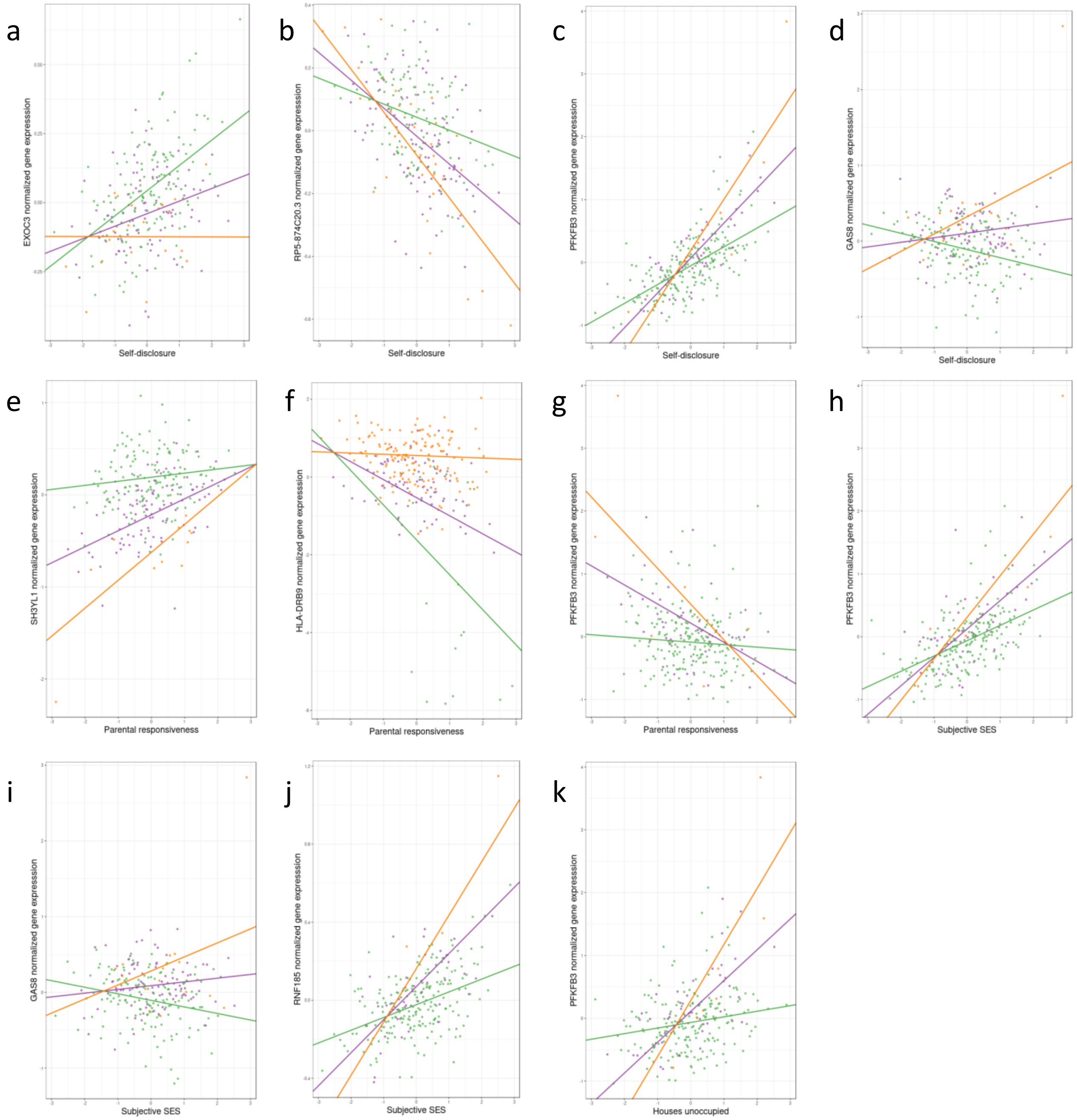


Fig. S11. Genetic variants interact with psychosocial environments to alter expression of genes linked to asthma and allergic disease. Scatterplots depict: a - Self-disclosure interacts with eQTL rs5865330:CT:C to alter expression of EXOC3 gene, b - Self-disclosure interacts with eQTL rs200494:T:G to alter expression of ZSCAN26 gene, c - Self-disclosure interacts with eQTL rs5015567:G:A to alter expression of PFKFB3, d - Self-disclosure interacts with eQTL rs12922757:G:A to alter expression of GAS8, e - parental responsiveness interacts with eQTL rs17713729:A:C to alter expression of SH3YL1, f - parental responsiveness interacts with eQTL rs9269774:G:A to alter expression of HLA-DRB9, g - parental responsiveness interacts with eQTL rs5015567 to alter expression of PFKFB3, h - socio-economic status interacts with eQTL rs5015567:G:A to alter expression of PFKFB3, i - socio-economic status interacts with eQTL rs12922757:G:A to alter expression of GAS8, j - socio-economic status interacts with eQTL rs36123367:AT:A to alter expression of RNF185, k – unoccupied houses interact with eQTL rs5015567:G:A to alter expression of PFKFB3.

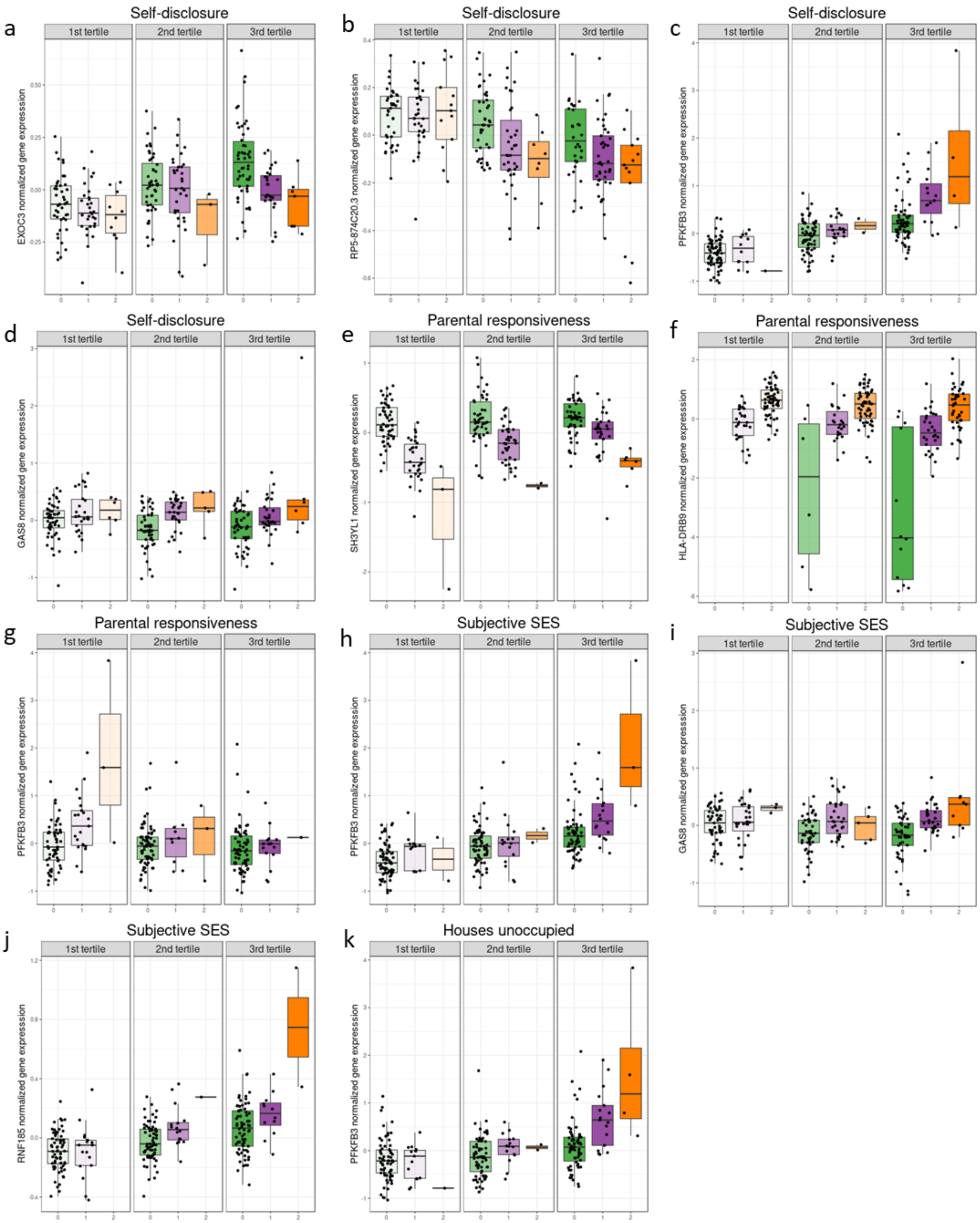

a
Self-disclosure
1st tertile
2nd tertile
3rd tertile
EXOC3 normalized gene expression
b
Self-disclosure
RP5-874C20.3 normalized gene expression
c
Self-disclosure
PFKFB3 normalized gene expression
d
Self-disclosure
GAS8 normalized gene expression
e
Parental responsiveness
SH3YL1 normalized gene expression
f
Parental responsiveness
HLA-DRB9 normalized gene expression
g
Parental responsiveness
PFKFB3 normalized gene expression
h
Subjective SES
PFKFB3 normalized gene expression
i
Subjective SES
GAS8 normalized gene expression
j
Subjective SES
RNF185 normalized gene expression
k
Houses unoccupied
PFKFB3 normalized gene expression

Fig. S12. Genetic variants interact with psychosocial environments to alter expression of genes linked to asthma and allergic disease. Boxplots depict: a - Self-disclosure interacts with eQTL rs5865330:CT:C to alter expression of EXOC3 gene, b - Self-disclosure interacts with eQTL rs200494:T:G to alter expression of ZSCAN26 gene, c - Self-disclosure interacts with eQTL rs5015567:G:A to alter expression of PFKFB3, d - Self-disclosure interacts with eQTL rs12922757:G:A to alter expression of GAS8, e - parental responsiveness interacts with eQTL rs17713729:A:C to alter expression of SH3YL1, f - parental responsiveness interacts with eQTL rs9269774:G:A to alter expression of HLA-DRB9, g - parental responsiveness interacts with eQTL rs5015567 to alter expression of PFKFB3, h - socio-economic status interacts with eQTL rs5015567:G:A to alter expression of PFKFB3, i - socio-economic status interacts with eQTL rs12922757:G:A to alter expression of GAS8, j - socio-economic status interacts with eQTL rs36123367:AT:A to alter expression of RNF185, k – unoccupied houses interact with eQTL rs5015567:G:A to alter expression of PFKFB3.

### 2. Supplementary tables:

All supplementary tables can be found in in file: Suppl_tables.xlsx

Table S1. Basic demographic information on 119 individuals for whom full data was available. Number reflects the count per each category, number in parenthesis reflects percentage.

Table S2. List of variables collected for current study. DD=daily diary, SD=Sleep diary, EAR=Coded from Electronically Activated Recorder, YR=Youth reported, PR=Parent reported, CD=Census data, GC=Glucocorticoid, SD=standard deviation, α=Chronbach's alpha measuring reliability as the average correlation between scale items, as a function of the number of items included in the scale.

Table S3. Correlations between the top 3 principal components of covariate matrix and individual covariates (Pearson product-moment correlations with numeric variables, polyserial correlations with bivariate variables; ns - correlation p-value>0.05).

Table S4. Differentially expressed genes associated with blood cell composition (10% FDR, DEG - differentially expressed genes, N - sample size).

Table S5. Differential gene expression results for all psychosocial variables (10% FDR, DEG - differentially expressed genes, N - sample size).

Table S6. Evaluation of transcriptional signatures derived using elastic net regression. For each variable we report Pearson's correlation coefficient, p-value, cross-validated percent variance explained, and sample size.

Table S7. Longitudinal replication of transcriptional signatures. For each variable we report Spearman's correlation coefficient between longitudinal change in observed variable and change in the transcriptional signature, p-value, coefficient of variation of longitudinal change in observed variable, and sample size.

### Supplementary files:

**File S1:** Detailed descriptions of methods for psychosocial data collection.

FileS1_Suppl_psych_measures_info.docx

**File S2:** Results of differential gene expression analysis with DESeq2.

FileS2_DESeq2_results.zip

**File S3:** GLMnet model weights for the transcriptional signatures. The transcriptional signatures are in columns, while the genes are in rows.

FileS3_glmnet_models.txt

**File S4**: Results of cis-eQTL mapping permutation pass with FastQTL: **a** - without correction for gene expression PCs, **b** - correcting for top 18 gene expression PCs (space-delimited files). Columns: 1 - Ensembl ID of the tested gene, 2 - Number of variants tested in cis for this gene, 3 - MLE of the shape1 parameter of the Beta distribution, 4 - MLE of the shape2 parameter of the Beta distribution, 5 - Dummy, 6 - ID of the best variant found for this molecular phenotypes (i.e. with the smallest p-value), 7 - Distance between the molecular phenotype - variant pair, 8 - The nominal p-value of association that quantifies how significant from 0, the regression coefficient is, 9 - The slope associated with the nominal p-value of association, 10 - A first permutation p-value directly obtained from the permutations with the direct method (corrected nominal p-value that accounts for the fact that multiple variants are tested per molecular phenotype), 11 - A second permutation p-value obtained via beta approximation [used to calculate FDR].

FileS4a_PC1-0.permutations.eQTL.txt.gz
FileS4b_PC1-18.permutations.eQTL.txt.gz

**File S5**: Results of cis-interaction-eQTL mapping (tab-delimited file). Columns: 1 - Ensembl ID of the tested gene, 2 - id of the tested variant, 3 - p-value of the GxE model intercept, 4 - p-value of the genotype dosage effect, 5 - p-value of the transcriptional signature effect, 6 - p-value of the interaction between genotype dosage and transcriptional signature effect, 7 - intercept, 8 - dosage effect, 9 - transcriptional signature effect, 10 - interaction between genotype dosage and transcriptional signature effect, 11 - intercept standard error (SE), 12 - genotype dosage effect SE, 13 - transcriptional signature effect SE, 14 - interaction between genotype dosage and transcriptional signature effect SE, 15 - q-value of the GxE model intercept, 16 - q-value of the genotype dosage effect, 17 - q-value of the transcriptional signature effect, 18 - q-value of the interaction between genotype dosage and transcriptional signature effect, 19 - tested transcriptional signature, 20 - permutation-corrected p-value of the interaction between genotype dosage and transcriptional signature effect, 21 - qvalue of the permutation-corrected p-value of the interaction between genotype dosage and transcriptional signature effect.

FileS5_GxE_0PCs_corrected.txt

**File S6**: Results of cis-interaction-eQTL mapping (tab-delimited file). Columns: 1 - Ensembl ID of the tested gene, 2 - id of the tested variant, 3 - p-value of the GxE model intercept, 4-7 - p-value of the imputed cell compositions effects, 8 - p-value of the genotype dosage effect, 9 - p-value of the transcriptional signature effect, 10 - p-value of the interaction between genotype dosage and transcriptional signature effect, 11 - intercept, 12-15 - imputed cell compositions effects, 16 - dosage effect, 17 - transcriptional signature effect, 18 - interaction between genotype dosage and transcriptional signature effect, 19 - intercept standard error (SE), 20-23 - the imputed cell compositions effects SEs, 24 - genotype dosage effect SE, 25 - transcriptional signature effect SE, 26 - interaction between genotype dosage and transcriptional signature effect SE, 27 - q-value of the GxE model intercept, 28-31 - q-values of the imputed cell compositions effects, 32 - q-value of the genotype dosage effect, 33 - q-value of the transcriptional signature effect, 34 - q-value of the interaction between genotype dosage and transcriptional signature effect, 35 - tested transcriptional signature, 36 - permutation-corrected p-value of the interaction between genotype dosage and transcriptional signature effect, 37 - qvalue of the permutation-corrected p-value of the interaction between genotype dosage and transcriptional signature effect.

FileS6_GxE_0PCs_cell-corrected.txt

**File S7:** Overlap of eGenes with psychosocial effects and significant PTWAS association results (5% FDR) for asthma and allergic disease. Columns: 1 - GWAS trait tested in PTWAS, 2 - Ensembl ID of the tested gene, 3 - Most significant tissue in PTWAS, 4 - multi-tissue PTWAS p-value; columns 5-25 - as in File S5.

FileS7_PTWAS-asthma-allergy_psychosocial_overlap.txt

**File S8:** Overlap of GxE interactions and significant PTWAS association results (5% FDR). Columns as in File S6.

FileS8_PTWAS_GxE_overlap.txt

### Supplementary text:

#### Participant recruitment methods

We used the following three recruitment methods, in order of most-used to least-used:

1. Dr. Secord, a co-I on the study, identifies patients who meet some of the basic criteria for the study who are then sent informational letters to their home address. The letter informs them of the basic information about the study and a telephone number to contact if interested.
2. A nurse from DMC Children's Hospital approaches patients and their guardians in asthma clinic or ER and informs them about the study. If interested, patient's guardian is later contacted to further inform them about the study and screen them for eligibility.
3. Through informational flyer, posted in local asthma clinic, distributed to local K-12 schools with basic information about the study and eligibility criteria, and study coordinator contact information.

Basic information provided to potential participants across the 3 methods informs them that this is a family asthma study looking at everyday life and how health is affected, that there is a home or office/lab visit focused on surveys, a brief interview and a 4 day at home period and a follow up where blood is collected.

#### Participant eligibility:

1. Child is between 10-15 years old.
2. Child has an asthma diagnosis of at least mild to persistent asthma.
3. At least one parent/guardian is willing to participate as well as the child.
4. Participating parent/guardian should be living with the child consistently for the last 6 months.
5. If unsure of which parent/guardian should participate, then the one that would have the most knowledge about health and day to day life.
6. If multiple children in the house have asthma and are in the study age range, the family can choose which child will participate based on who would be most agreeable to the various measures taken throughout the study.

#### Identification of gene expression confounders

There are many sources of technical and biological noise in RNA-seq data. To accurately estimate gene expression differences between meaningful groups, one must account for additional sources of gene expression variation that are not of interest. To investigate the major sources of variation in the RNA-seq dataset we performed principal component analysis (PCA) on normalized RNA-seq data and investigated 1) the correlation between first 15 PCs and each of the covariates (data not shown), and 2) proportion of variance explained by each covariate overall (Fig. S2a), and per each gene (Fig. S2b). The following covariates were significantly correlated with at least one PC: library preparation batch (confounded with library sequencing batch), RNA quality score (RIN), laboratory where the RNA sample was extracted, age, sex, height, weight and ancestry (measured as first three principal components on the genotype matrix). These variables were also correlated with each other, therefore we used PCA to calculate the PCs of all these covariates that explained >99.5% variance and incorporated them in the gene expression analyses as described in the methods.

##### Blood composition has a profound effect on gene expression

Blood composition is expected to have a strong effect on overall blood gene expression as blood is a heterogeneous tissue and different cell types may contribute different transcripts to the overall gene expression. The results of differential gene expression analysis with DESeq2 using likelihood ratio test (LRT) are displayed in Table S3. As expected, the number of differentially expressed genes is larger for cell types that constitute a bigger fraction of the cell pool. Accordingly, we did not find genes differentially expressed for basophils, a cell type that constitutes less than 1% of total leukocytes. Based on these results, cell composition was accounted for in differential gene expression analyses of psychosocial variables and other analyses that distinguished between the two types of effects (mediation analysis).

##### Differential gene expression analysis reveals similarities in gene expression among positive or negative psychosocial experiences

We ran principal component analysis of z-scores from differential gene expression analysis (see methods). PC1 explains 24% of variance, with subjective SES at one extreme and parental depression on the opposite extreme along this PC. Emotionality variables align from negative (top) to positive (bottom, Fig. S5) along PC2, which explains 16% of variance. Social relationship variables also tend to follow the positive-to-negative gradient along PC2. These results demonstrate that negative psychosocial experiences tend to be associated with similar gene expression changes, which can be distinguished from gene expression changes associated with positive experiences.

##### Overlap of transcriptional signatures

We investigated whether transcriptional signatures for different variables were correlated with each other, which would suggest shared transcriptional pathways for these phenotypes and environments (Fig. S6a). Transcriptional signatures of the socioeconomic measures showed strong overlap: specifically, houses rated ≥fair, unoccupied houses, parental income were all significantly correlated with one another, and parental income and subjective SES was correlated with each of those (but not one another). There was also some overlap in the social relationships category, with transcriptional signatures of daily youth-reported self-disclosure, daily youth-reported responsiveness, and objective parental responsiveness all significantly intercorrelated. However, we also saw correlations crossing all three variable categories. For example, socioeconomic status was significantly correlated with objective parental responsiveness, family conflict, and self-disclosure; objective negative affect was correlated with self-disclosure, familial conflict, maternal responsiveness and houses rated ≥fair.

We were able to identify transcriptional signatures that explained at least 1% of variance for 4 of the 5 blood composition variables and 3 of the 9 glucocorticoid variables. Models predicting blood composition performed best with 47-150 genes, and models predicting glucocorticoid variables contained 17-100 genes. Notably, the transcriptional signatures explained a far greater percent of variation in blood composition than other categories of variables. Transcriptional signatures within each category were highly correlated, showing greater overlap within all glucocorticoid response variables and all blood composition variables (except for eosinophils, which were uncorrelated with all other blood composition signatures) than seen in asthma severity or social relationship categories. When we correlated glucocorticoid and blood composition transcriptional signatures with transcriptional signatures for psychosocial experiences and asthma severity, we observed some overlap for glucocorticoid response: specifically, stimulated levels of IL-5 were associated with percent-predicted FEV1 (r= -.47, p<.001), unoccupied houses (r= -.19, p=.04), and youth reported parent-child conflict (r=.23, p=.01). There was much more overlap in blood composition transcriptional signatures with asthma severity and psychosocial experience signatures. For example, transcriptional signature of monocytes was correlated with a large number of transcriptional signatures of both asthma and psychosocial experiences, including nightly asthma symptoms (r=.19, p=.04), and FEV (r=.37, p<.001), self-disclosure (r=.46, p<.001), unoccupied houses (r=.29, p=.002), subjective SES (r=.37, p<.001), and objective negative affect (r=.41, p<.001). Lymphocytes showed a very similar pattern of correlations to neutrophils, but in the opposite direction.

#### Targeted vs. unbiased approach to identifying effects of psychosocial experiences on gene expression

Previous research analyzing blood gene expression developed the Conserved Transcriptional Response to Adversity (CTRA[1]), which proposes expression of 53 immune genes as a composite indicator of immune system response to adverse social environments (reviewed in[2]). We compared the performance of our unbiased prediction model with a model limited to the 53 CTRA genes (Fig. S7). The CTRA-based model performs better at predicting the fraction of two of the five blood cell types. This may reflect differential expression of CTRA genes in some cell types. CTRA-based and unbiased models perform similarly on psychosocial and neighborhood measures, with 9/16 traits (56%) predicted more accurately by the unbiased model. CTRA-based model performs poorly at predicting asthma and glucocorticoid phenotypes, and is outperformed by the unbiased model in 10/14 traits (71%) which is expected because it was not originally developed to capture these phenotypes. In summary, limiting the scope of the prediction model to preselected immune genes does not greatly diminish performance in predicting blood and psychosocial phenotypes, but the CTRA-based model does not appropriately reflect transcriptomic signatures of asthma and glucocorticoid phenotypes. This result suggests that genes outside the CTRA subset are important to many asthma and some psychosocial phenotypes.

#### Transcriptional signatures aid in denoising the data

We assume that the observed variable has high levels of noise and the measured value does not reflect the true biological effect. Therefore, we use the predicted values for all participants, including those for whom the variable was directly measured (denoising). The predicted values are imputed based on the generalized linear models with penalized maximum likelihood built using *glmnet* for each variable separately, according to the general model: Phenotype or Environment = $intercept + \beta_1 \mathrm{E}(\mathrm{gene}_1) + \beta_2 \mathrm{E}(\mathrm{gene}_2) + \ldots + \beta_n \mathrm{E}(\mathrm{gene}_n)$, where $\mathrm{E}(\mathrm{gene}_{\mathrm{n}})$ is normalized expression of gene n, $\beta_n$ is its estimated coefficient, and n is minimized via penalized maximum likelihood with elastic-net mixing parameter α set to 0.1 (0 representing ridge regression, 1 representing lasso regression).

The best fit for the model will be used to predict, for both observed and unobserved samples, the biological impact on gene expression of the relevant variable. We do this because the measurement error for the observed variables will also not be uniform as some individuals may not respond accurately or truthfully (e.g., to self-disclosure questions) to all questions. If the objective was to estimate the measurement including its biases and errors, the "technical" variance for the observed variable would be smaller if we were to use the observed values. However, we use the denoised/imputed values, where the "biological" variance, or the error between the fitted value of the signature with respect to the true unobserved biological impact, would be the same for both the measured and non-measured individuals. Overall, our procedure should not create any bias but it will decrease the variability of the imputed/denoised variables thus reducing the chance of false positive GxE.

#### Replication of GxE effects

To validate the observed GxE effects on gene expression we explored the overlap between the GxE genes and previously published datasets that measured interactions with different environments. We found that the majority of our GxE genes replicated in other datasets of GxE in gene expression ($p<0.05$). Of our genes with GxE, 2 were found to have GxE in response to influenza[3], 59 were found to have GxE in response to a variety of chemical treatments[4], 75 were found to have GxE in response to a variety of M. tuberculosis[5], 87 were found to have GxE in response to rhinovirus[6], 99 genes were found to have GxE in response to pathogens[7], and 117 had GxE with cell type fraction in whole blood[8].
