## Supplementary material for "Psychosocial experiences modulate asthma-associated genes through gene-environment interactions": File S5

Link to File S5:

[http://genome.grid.wayne.edu/ALOFT/Supp/FileS5_GxE_0PCs_corrected.txt.gz](https://www.google.com/url?q=http://genome.grid.wayne.edu/ALOFT/Supp/FileS5_GxE_0PCs_corrected.txt.gz&sa=D&source=hangouts&ust=1576366176636000&usg=AFQjCNGhPW4iIRWQ1_6YrtGcF9E-sR3IuA" \t "_blank)
