## Supplementary material for "Psychosocial experiences modulate asthma-associated genes through gene-environment interactions": File S1

### MacArthur Status Ladder (PR Subjective socioeconomic status)

**From:** Adler, N. E., Epel, E. S., Castellazzo, G., Ickovics, & J. R. (2000).  Relationship of subjective and objective social status with psychological and physiological functioning: Preliminary data in healthy, White women.  Health Psychology, 19, 586-592.

**Composite Formation and Variable Names:**

| Socio-Economic Status Ladder |  | psesl |
| --- | --- | --- |

**Response Scales:**

Range: 1 (*Lowest status*) - 10 (*Best status*)

**Data Note:**

The ladder coding was adjusted so lower scores represent lower status and higher scores represent better status.

### Neighborhood Stress Inventory (PR Neighborhood stress)

**From:** Ewart, C.K. & Suchday, S. (2002). Discovering how urban poverty and violence affect health: Development and validation of a neighborhood stress index. *Health Psychology, 21,* 254-262.

**Composite Formation:**

| **Neighborhood Disorder:** | |  |  |
| --- | --- | --- | --- |
|  | I saw people dealing drugs near my home |  | pnsi1 |
|  | I saw strangers who were drunk or high hanging out near my home |  | pnsi2 |
|  | I heard adults arguing loudly on my street |  | pnsi3 |
|  | I heard neighbors complaining about crime in our neighborhood |  | pnsi4 |
|  | Someone I knew was arrested or went to jail |  | pnsi5 |
|  | I saw or heard about a shooting gallery near my home |  | pnsi6 |
|  | People in the neighborhood complained about being harassed by police |  | pnsi7 |
|  | There was a gang fight near my phone |  | pnsi8 |
|  | About how many of your neighbors do you think received food stamps in the past year? |  | pnsi9 |
|  | About how many of your neighbors do you think received food stamps in the past year? |  | pnsi10 |
|  | I saw cars speeding or driving dangerously on my street |  | pnsi18 |
|  | **Mean Score** |  | **pnsnd** |
| **Total Mean Score:** at least 15 of 18 items | |  | **pnsi** |
| **Neighborhood Violence:** | |  |  |
|  | A family member was attacked or beaten |  | pnsi11 |
|  | A family member was stabbed or shot |  | pnsi12 |
|  | A family member was stopped and questioned by the police |  | pnsi13 |
|  | A friend was robbed or mugged |  | pnsi14 |
|  | A friend was stabbed or shot |  | pnsi15 |
|  | Someone threatened to hurt a member of my family |  | pnsi16 |
|  | A family member was robbed or mugged |  | pnsi17 |
|  | **Mean Score** |  | **pnsev** |

**Response Scales:**

pnsi1-pnsi8, pnsi11-pnsi18: 1 (*Never*), 2 (*Once*), 3 (*A few times*), 4 (*Often*)

pnsi9-pnsi10: 1 (*None*), 2(*Some*), 3(*About half*), 4 (*Most*)

**Data Note:**

Higher scores reflect greater neighborhood stress

pnsi18 was added 9/14/12

### Child Daily Perceived “Other” Responsiveness to Disclosure (DD Perceived responsiveness)

**Adapted From:** Laurenceau, J., Barrett, L. F., & Pietromonaco, P. R. (1998). Intimacy as a process: The importance of self-disclosure and responsiveness in interpersonal exchanges. Journal of Personality and Social Psychology, 74, 1238-1251.

**Composite Formation and Variable Name:**

| **Child Perceived Responsiveness** | |  |  |
| --- | --- | --- | --- |
| *After talking with this person, did you feel…* | |  |  |
|  | That the person really listened to what you were saying |  | cdres1 |
|  | That the person was responsive to what you were saying |  | cdres2 |
|  | Accepted by that person |  | cdres3 |
|  | **Mean Score:** mean of at least 2 of the 3 items |  | **cdres** |

**Response Scale:**

Range: 1 (*Not At All*) – 5 (*Extremely*)

### Child Daily Diary Self-Disclosure (DD Self-disclosure)

**Adapted From:** Laurenceau, J., Barrett, L. F., & Pietromonaco, P. R. (1998). Intimacy as a process: The importance of self-disclosure and responsiveness in interpersonal exchanges. Journal of Personality and Social Psychology, 74, 1238-1251.

| **Child Daily Disclosure** | |  |  |
| --- | --- | --- | --- |
| *I talked about…* | |  |  |
|  | Facts and information | cddi1 |  |
|  | Your thoughts | cddi2 | cddis2 |
|  | Your feelings | cddi3 | cddis3 |
|  | **Mean:** at least 2 of the 3 items | **cdds** | **cddstf** |

**Response Scale**:
Range: 1 (*Not At All*) – 5 (*Extremely*)

### Parental Warmth (YR Parental warmth)

**From:** Schaefer, E.S. (1965). Children’s Reports of Parental Behavior: An Inventory. *Child Development, 36*(2), 413-424

**Composite Formation & Variable Name:**

| Mother – Parental Warmth – low child baseline | |  |  |
| --- | --- | --- | --- |
|  | My mother (or female guardian) always listens to my ideas and opinions. |  | cpwm1 |
|  | My mother (or female guardian) often talks about the good things I do. |  | cpwm2 |
|  | My mother (or female guardian) makes me feel better when I am upset. |  | cpwm3 |
|  | My mother (or female guardian) gives me a lot of care and attention. |  | cpwm4 |
|  | My mother (or female guardian) seems proud of the things I do. |  | cpwm5 |
|  | My mother (or female guardian) says I make her happy. |  | cpwm6 |
|  | My mother (or female guardian) is happy to see me when I come home. |  | cpwm7 |
|  | My mother (or female guardian) shows his love for me. |  | cpwm8 |
|  | My mother (or female guardian) almost always talks to me with a warm and friendly voice. |  | cpwm9 |
|  | My mother (or female guardian) is always thinking of things that will please me. |  | cpwm10 |
|  | My mother (or female guardian) often praises me. |  | cpwm11 |
|  | My mother (or female guardian) makes me feel better after talking over my worries with her. |  | cpwm12 |
|  | My mother (or female guardian) smiles at me very often |  | cpwm13 |
|  | My mother (or female guardian) tells me how much she loves me. |  | cpwm14 |
|  | My mother (or female guardian) cheers me up when I am sad. |  | cpwm15 |
|  | My mother (or female guardian) is very interested in what I have to say. |  | cpwm16 |
|  | My mother (or female guardian) understands how I see things. |  | cpwm17 |
|  | My mother (or female guardian) is a friend. |  | cpwmd18 |
|  | My mother (or female guardian) seems to be thinking of me. |  | cpwm19 |
|  | My mother (or female guardian) shows me that she loves me. |  | cpwm20 |
|  | There is a feeling of warmth between my mother (or female guardian) and me. |  | cpwm21 |
|  | There is a feeling of togetherness between my mother (or female guardian) and me. |  | cpwm22 |
|  | My mother (or female guardian) and I show affection toward each other. |  | cpwm23 |
|  | **Mean Score**: at least 18 of the 23 items |  | **cpwml** |
|  | Reverse-scored composite |  | **cpwm** |

**Response Scales:**

Range: 1 (*Agree*), 2 (*Somewhat Agree*), 3 (*Disagree*)

**Data Note:**

High Child Baseline - higher scores reflect greater parental warmth

### Child Daily Diary Family Interaction Scales (DD Marital conflict and DD Marital affection)

**Adapted from:** the *Youth Everyday Social Interactions and Mood Scales (YES-I-AM)^1,2^* and the Child Home Data Questionnaire^3^

^1^ Repetti, R. L. (1996). The Effects of Perceived Daily Social and Academic Failure Experiences on School-Age Children’s Subsequent Interactions with Parents. *Child Development,* 67(4), 1467-1482.

^2^ Repetti, R.L., & Polina, S.L. (1994). *Development of child daily-report measures: The Youth Everyday Social Interaction and Mood (YES I AM) Scales*. Unpublished manuscript.

^3^ Margolin, G. (1990). *Child Home Data Questionnaire.* Unpublished Manuscript. Los Angeles: University of Southern California.

**Composite Formation and Variable Name:**

| *Marital Interactions (Child Home Data Questionnaire):* | |  |  |
| --- | --- | --- | --- |
| **Child-reported Marital Conflict** | |  |  |
|  | My mom and dad seemed angry with each other today |  | cdpin1 |
|  | My mom and dad argued today |  | cdpin2 |
|  | **Total Score** |  | **cdmcfl** |
| **Child-reported Aversive Behavior with Mother** | |  |  |
|  | My mom and dad kissed or hugged today |  | cdpin3 |
|  | **Item as reported in database** |  | **cdmaff** |

**Data Note:**

All family interaction scale scores are means unless the scale consists of a single item (at least 2 of 3 items).

Responses are to Mom or female guardian/ Dad or male guardian

**Response Scales:**
Range: 1 (*Not At All*), 2 (*Some*), 3 (*A Lot*)

### Parental Environment Questionnaire (PEQ) (YR Parent-child conflict)

**From:** Elkins, I.J., McGue, M., & Iacono, W.G. (1997). Genetic and environmental influences on parent-son relationships: evidence for increasing genetic influence during adolescence. *Developmental Psychology, 33,* 351-363.

**Composite Formation and Variable Name:**

| **Conflict - Mother** | |  |  |
| --- | --- | --- | --- |
|  | My mother (or female guardian) often criticizes me |  | cpeqm1 |
|  | Before I finish saying something, my mother (or female guardian) often interrupts me |  | cpeqm2 |
|  | My mother (or female guardian) often irritates me |  | cpeqm3 |
|  | There are often misunderstandings between my mother (or female guardian) and me |  | cpeqm4 |
|  | I treat others with more respect than I treat my mother (or female guardian) |  | cpeqm5 |
|  | My mother (or female guardian) often hurts my feelings |  | cpeqm6 |
|  | My mother (or female guardian) does not trust me to make my own decisions |  | cpeqm7 |
|  | My mother (or female guardian) and I often get into arguments |  | cpeqm8 |
|  | I often seem to anger or annoy my mother (or female guardian) |  | cpeqm9 |
|  | My mother (or female guardian) often loses her temper with me |  | cpeqm10 |
|  | My mother (or female guardian) sometimes hits me in anger |  | cpeqm11 |
|  | Once in a while I have been really scared of my mother (or female guardian) |  | cpeqm12 |
|  | **Mean Score:** mean of at least 8 of the 12 items |  | **cpeqcm** |

**Response Scales:**

Range: 1 (*Definitely False*), 2 (*Probably False*), 3 (*Probably True*), 4 (*Definitely True*)

Reverse coded items designated by ‘r’

**Data Note:**

PEQ is coded so higher scores reflect higher conflict or higher involvement

### Parental Environment Questionnaire (PEQ) (PR Parent-child conflict)

**From:** Elkins, I.J., McGue, M., & Iacono, W.G. (1997). Genetic and environmental influences on parent-son relationships: evidence for increasing genetic influence during adolescence. *Developmental Psychology, 33,* 351-363.

**Composite Formation and Variable Names:**

| **Conflict** | |  |  |
| --- | --- | --- | --- |
|  | I often lose my temper with my child |  | ppeq1r |
|  | I often have misunderstandings with my child |  | ppeq2r |
|  | My child and I often argue |  | ppeq3r |
|  | I often criticize my child |  | ppeq4r |
|  | My child often angers or annoys me |  | ppeq5r |
|  | I often hurt my child’s feelings |  | ppeq6r |
|  | I often irritate my child |  | ppeq7r |
|  | I sometimes hit my child in anger |  | ppeq8r |
|  | My child has been really scared of me |  | ppeq9 |
|  | I often interrupt my child |  | ppeq10r |
|  | My child respects others more than me |  | ppeq11r |
|  | I often do not trust my child’s decision |  | ppeq12r |
|  | **Mean Score:** at least 9 of 12 items |  | **ppeqcf** |

**Response Scales:**

Range: 1 (*Definitely true*), 2 (*Probably True*), 3 (*Probably False*), 4 (*Definitely false*)

**Data Note:**

PEQ is coded so that high scores reflect higher conflict or higher involvement (‘r’ indicates reverse coded item)

ALOFT only uses two of the possible subscales for the questionnaire

**Center for Epidemiological Studies Depression Scale (CES-D) (PR Parental depressive symptoms)**

**From:** Radloff, L. S. (1977). The CES-D scale: A self-report depression scale for research in the general population. *Applied Psychological Measurement, 1*, 385-401.

**Composite Formation and Variable Names:**

| *During the past week:* | |  |  |
| --- | --- | --- | --- |
|  | I was bothered by things that usually don’t bother me |  | pcesd1r |
|  | I did not feel like eating; my appetite was poor |  | pcesd2r |
|  | I felt that I could not shake off the blues even with help from my family or friends |  | pcesd3r |
|  | I felt like I was just as good as other people |  | pcesd4rr |
|  | I had trouble keeping my mind on what I was doing |  | pcesd5r |
|  | I was depressed |  | pcesd6r |
|  | I felt that everything I did was an effort |  | pcesd7r |
|  | I felt hopeful about the future |  | pcesd8rr |
|  | I thought my life had been a failure |  | pcesd9r |
|  | I felt fearful |  | pcesd10r |
|  | My sleep was restless |  | pcesd11r |
|  | I was happy |  | pcesd12rr |
|  | I talked less than usual |  | pcesd13r |
|  | I felt lonely |  | pcesd14r |
|  | People were unfriendly |  | pcesd15r |
|  | I enjoyed life |  | pcesd16rr |
|  | I had crying spells |  | pcesd17r |
|  | I felt sad |  | pcesd18r |
|  | I felt that people disliked me |  | pcesd19r |
|  | I could not get going |  | pcesd20r |
|  | **Total Sum:** |  | **pcesdts** |
|  | **Mean Score:** |  | **pcesdm** |
|  | **Mean Score:** at least 16 of 20 items |  | **pcesdts16** |

**Response Scale:**

Range: 0 (*Rarely or none of the time (less than 1 day)*), 2 (*A Little Amount of the Time*), 3(*A Moderate Amount of the Time*), 4 (*Most of or all of the time (5-7 days)*)

**Data Note:**

Items were recoded to start with 0 instead of 1 indicated with an ‘r’

Reverse coded items indicated by an additional ‘r’

Higher scores reflect more depressive symptoms during the last week

### Risky Families Questionnaire (YR Risky family environment)

**From:** Felitti, V. J., Anda, R. F., Nordenberg, D., Williamson, D. F., Apitz, A. M., Edwards, V., et al. (1998). Relationship of childhood abuse and household dysfunction to many of the leading causes of death in adults. *American Journal of Preventative Medicine, 14*, 245-258.

Taylor, S. E., Lerner, J. S., Sage, R. M., Lehman, B. J., & Seeman, T. E. (2004). Early environment, emotions, responses to stress, and health. *Journal of Personality, 72*, 1365-1393.

**Composite Formation and Variable Name:**

| How often does a parent or other adult in your household make you feel that you are loved, supported, and cared for? |  | crfq1*r* |
| --- | --- | --- |
| How often does a parent or other adult in your household swear at you, insult you, put you down, or act in a way that makes you feel threatened? |  | crfq2 |
| How often does a parent or other adult in your household express physical affection to you, such as hugging, or other physical gestures of warmth and affection? |  | crfq3*r* |
| How often does a parent or other adult in your household push, grab, shove, or slap you? |  | crfq4 |
| Do you live with anyone who is a problem drinker or alcoholic, or used street drugs? |  | crfq5 |
| Would you say that the household you live in is well organized and well managed? |  | crfq6*r* |
| How often would you say that a parent or other adult in the household behaves violently toward a family member or visitor in your home? |  | crfq7 |
| How often would you say there is quarreling, arguing, or shouting between your parents? |  | crfq8 |
| How often would you say there is quarreling, arguing, or shouting between a parent and you? |  | crfq9 |
| How often would you say there is quarreling, arguing, or shouting between a parent and one of your siblings? |  | crfq10*r* |
| How often would you say there is quarreling, arguing, or shouting between your sibling(s) and you? |  | crfq11*r* |
| Would you say that your household is chaotic and disorganized? |  | crfq12 |
| How often would you say you are neglected, that is, left ln your own to fend for yourself? |  | crfq13 |
| **Mean Score**: mean of at least 7 items |  | **criskf** |

**Response Scales:**

Range: 1 (*Never*), 2 (*Rarely*), 3 (*Sometimes*), 4 (*Often*), 5 (*Very often*)

Reverse coded items designated by ‘*r*’

Not applicable items recoded as missing values: crfq10*r*, crfq11*r*

**Data Note:**

Higher scores reflect higher risk within the family

### Child Daily Diary Mood Scales and Clusters (DD Positive affect and DD Negative affect)

**From:** Cohen et al. (2006). Positive Emotional Style Predicts Resistance to Illness after Experimental Exposure to Rhinovirus or Influenza A Virus. *Psychosomatic Medicine,68*, 809–815.

Doyle, W.J., Gentile, D.A., & Cohen, S. (2006). Emotional style, nasal cytokines, and illness expression after experimental rhinovirus exposure. *Brain, Behavior, and Immunity, 20*, 175–181.

**Composite Formation and Variable Name:**

| **Child Positive Mood** (at least 6 of 8 items) | |  | **cdposm** |
| --- | --- | --- | --- |
|  | 1. Lively |  | (*cdfl1*) |
|  | 3. Happy |  | (c*dfl3*) |
|  | 5. At ease |  | (c*dfl5*) |
|  | 8. Full of energy |  | (c*dfl8*) |
|  | 10. Cheerful |  | (c*dfl10*) |
|  | 12. Calm |  | (c*dfl12*) |
|  | 14. proud^b^ |  | (c*dfl14*) |
|  | 15. loved^b^ |  | (c*dfl15*) |
| **Child Negative Mood** (at least 4 of 6 items) | | **cdnegm** | **cdngm2** |
|  | 2. Sad | (*cdfl2*) | (*cdfl2*) |
|  | 4. On edge |  | (*cdfl4*) |
|  | 6. Hostile |  | (*cdfl6*) |
|  | 7. Mean^a^ | (*cdfl7*) | (*cdfl7*) |
|  | 9. Unhappy | (*cdfl9*) | (*cdfl9*) |
|  | 11. Tense | (*cdfl11*) | (*cdfl11*) |
|  | 13. Angry | (*cdfl13*) | (*cdfl13*) |
|  | 16. Worried^a^ | (*cdfl16*) | (*cdfl16*) |

**Responses Scale**:

Range: 1 (*Completely inaccurate*) – 4 (*Completely accurate*)

**Data Note:**

All scale scores are means unless otherwise stated or the scale consists of a single item.

*Mood Scale items differed slightly between parent and child diaries. Some items were added to be more age appropriate. Two items were also added from Repetti’s YES I AM mood scales:

^a^ Items reworded from adult mood scale to aid in child comprehension (added)

‘Mean’ reworded from adult version’s ‘hostile’

‘Worried’ reworded from adult version’s ‘on edge’

^b^ Items added from Repetti’s YES I AM mood scales and do not appear in Positive Mood clusters

‘Proud’

‘Loved’

### Child Depression Inventory (CDI) (YR Depressive symptoms)

**From:** Kovacs, M. (1992). *Children's Depression Inventory: Manual*. New York: Multi-Health Systems.

**Composite Formation and Variable Name:**

| *Pick out the sentences that describe your feelings and thoughts in the past two weeks* | |  |  |
| --- | --- | --- | --- |
|  | 1 (*I am sad once in a while*), 2 (*I am sad many times*), 3 (*I am sad all the time*) |  | ccdi1 |
|  | 1 (*Nothing will ever work out for me*), 2 (*I am not sure if things will work out for me*), 3 (*Things will work out right for me*) |  | ccdi2r |
|  | 1 (*I do most things right*), 2 (*I do many things wrong*), 3 (*I do everything wron*g) |  | ccdi3 |
|  | 1 (*I have fun in many things*), 2 (*I have fun in some things*), 3 (*Nothing is fun at all*) |  | ccdi4 |
|  | 1 (*I am bad all the time*), 2 (*I am bad many times*), 3 (*I am bad once in a while*) |  | ccdi5r |
|  | 1 (*I think about bad things happening to me once in a while*), 2 (*I worry that bad things will happen to me*), 3 (*I am sure that terrible things will happen to me*) |  | ccdi6 |
|  | 1 (*I hate myself*), 2 (*I do not like myself*), 3 (*I like myself*) |  | ccdi7r |
|  | 1 (*All bad things are my fault*), 2 (*Many bad things are my fault*), 3 (*Bad things are not usually my fault*) |  | ccdi8r |
|  | 1 (*I do not think about killing myself*), 2 (*I think about killing myself but I would not do it*), 3 (*I want to kill myself*) |  | ccdi9 |
|  | 1 (*I feel like crying everyday*), 2 (*I often feel like crying*), 3 (*I feel like crying once in a while*) |  | ccdi10r |
|  | 1 (*Things bother me all the time*), 2 (*Things often bother me*), 3 (*Things bother me once in a while*) |  | ccdi11r |
|  | 1 (*I like being with people*), 2 (*I do not like being with people many times*), 3 (*I do not want to be with people at all*) |  | ccdi12 |
|  | 1 (*I cannot make up my mind about things*), 2 (*It is hard to make up my mind about things*), 3 (*I make up my mind about things easily*) |  | ccdi13r |
|  | 1 (*I look O.K.*), 2 (*There are some bad things about my looks*), 3 (*I look ugly*) |  | ccdi14 |
|  | 1 (*I have to push myself all the time to do my schoolwork*), 2 (*I have to push myself many times to do my schoolwork*), 3 (*Doing schoolwork is not a big problem*) |  | ccdi15r |
|  | 1 (*I have trouble sleeping every night*), 2 (*I have trouble sleeping many nights*), 3 (*I sleep pretty well*) |  | ccdi16r |
|  | 1 (*I am tired once in a while*), 2 (*I am tired many days*), 3 (*I am tired all the time*) |  | ccdi17 |
|  | 1 (*Most days I do not feel like eating*), 2 (*Many days I do not feel like eating*), 3 (*I eat pretty well*) |  | ccdi18r |
|  | 1 (*I do not worry about aches and pains*), 2 (*I worry about aches and pains many times*), 3 (*I worry about aches and pains all the time*) |  | ccdi19 |
|  | 1 (*I do not feel alone*), 2 (*I feel alone often*), 3 (*I feel alone all the time*) |  | ccdi20 |
|  | 1 (*I never have fun at school*), 2 (*I have fun at school only once in a while*), 3 (*I have fun at school many times*) |  | ccdi21r |
|  | 1 (*I have plenty of friends*), 2 (*I have some friends but I wish I had more*), 3 (*I do not have any friends*) |  | ccdi22 |
|  | 1 (*My schoolwork is alright*), 2 (*My schoolwork is not as good as before*), 3 (*I do very badly in subjects I used to be good in*) |  | ccdi23 |
|  | 1 (*I can never be as good as other kids*), 2 (*I can be as good as other kids if I want to be*), 3 (*I am just as good as other kids*) |  | ccdi24r |
|  | 1 (*Nobody really loves me*), 2 (*I am not sure if anybody loves me*), 3 (*I am sure that somebody loves me*) |  | ccdi25r |
|  | 1 (*I usually do that I am told*), 2 (*I do not do what I am told most times*), 3 (*I never do what I am told*) |  | ccdi26 |
|  | 1 (*I get along with people*), 2 (*I get into fights many times*), 3 (*I get into fights all the time*) |  | ccdi27 |
|  | **Total Score** is sum of all 27 items |  | **ccdiTS** |
|  | **Mean Score**: at least 24 of the 27 items |  | **ccdim** |

**Response Scales:**

Recoded all values from (1 – 3) range to (0 – 2) range

Reverse coded items designated by ‘r’

**Data Note:**

Higher scores reflect greater depressive symptoms
